# Rapid Reconstruction of Time-varying Gene Regulatory Networks with Limited Main Memory

**DOI:** 10.1101/755249

**Authors:** Saptarshi Pyne, Ashish Anand

## Abstract

Reconstruction of time-varying gene regulatory networks underlying a time-series gene expression data is a fundamental challenge in the computational systems biology. The challenge increases multi-fold if the target networks need to be constructed for hundreds to thousands of genes. There have been constant efforts to design an algorithm that can perform the reconstruction task correctly as well as can scale efficiently (with respect to both time and memory) to such a large number of genes. However, the existing algorithms either do not offer time-efficiency, or they offer it at other costs – memory-inefficiency or imposition of a constraint, known as the ‘smoothly time-varying assumption’. In this paper, two novel algorithms – ‘an algorithm for reconstructing Time-varying Gene regulatory networks with Shortlisted candidate regulators - which is Light on memory’ (*TGS-Lite*) and ‘TGS-Lite Plus’ (*TGS-Lite+*) – are proposed that are time-efficient, memory-efficient and do not impose the smoothly time-varying assumption. Additionally, they offer state-of-the-art reconstruction correctness as demonstrated with three benchmark datasets.

**Source Code:** https://github.com/sap01/TGS-Lite-supplem/tree/master/sourcecode

## 1 Introduction

PROTEINS perform a multitude of functions in numerous biological processes. The expression of every protein depends on various factors; one of them is the expression of the gene that encodes that protein. The expression of a gene, in turn, can be regulated by that of one or more genes, which are known as the regulators of the former gene (the regulatee). These regulator-regulatee relationships are represented as a directed network, known as the Gene Regulatory Network (GRN) [1]. In a GRN, the nodes represent the genes and the edges represent their relationships. Since all the regulators of a regulatee may not remain active all the time, the edge relationships (hereafter, structure) of a GRN may vary with time. Discovering how the GRN structure underlying a biological process varies with time is considered to be a fundamental question in the systems biology ([1], [2]). Being able to answer this question helps in understanding the underlying mechanisms of the concerned biological process, such as developmental programs and pathogenesis, at a molecular level.

Since the structural changes are reflected in the expressions of the genes, time-series gene expressions are collected in an attempt to reverse-engineer (reconstruct) how the structure of the underlying GRN varies with time. For high-throughput time series gene expression datasets, it becomes infeasible to conduct the reconstruction task manually; instead, computational algorithms are employed for that purpose. Such algorithms are called time-varying GRN reconstruction algorithms ([3], [4]).

There exists a bevy of algorithms ([4], [5], [6], [7], [8], [9], [10], [11], [12], [13], [14]) that can be used to reconstruct time-varying GRNs from time-series gene expression datasets. These algorithms can be broadly divided into two categories: ones that are time-intensive ([4], [5], [6], [7], [8], [11]) and the others that achieve time-efficiency by imposing a constraint, known as the ‘smoothly time-varying assumption’ ([9], [10], [12], [13]), or by compromising on memory-efficiency ([14]). A comparative study of a subset of these algorithms ([8], [9], [14]) is conducted by Pyne et al. [14] against three benchmark datasets. The study demonstrates that the time-intensive algorithms and the memory-inefficient algorithms tend to reconstruct GRNs more correctly than the algorithms with the smoothly time-varying assumption. More specifically, a time-intensive algorithm, namely *ARTIVA* [8], most consistently produces the lowest numbers of false positives. On the other hand, a fast but memory-inefficient algorithm, namely *TGS* [14], most consistently produces the highest numbers of true positives. Another fast but memory-inefficient algorithm, namely *TGS+* [14], finds the desired balance: it consistently achieves a comparable number of false positives to those of *ARTIVA* and a high number of true positives comparable to those of *TGS*. Nevertheless, the Achilles’ heel of both *TGS* and *TGS+* is their main memory (hereafter, simply ‘memory’) requirements, which grow exponentially with the number of genes in the given dataset. As Jahnsson et al. aptly put it, “The algorithm can always be given more time; however, if it exceeds the available memory resources, nothing can be done to solve the instance” [15].

In this paper, two novel algorithms – *TGS-Lite* and *TGS-Lite+* – are proposed. None of them imposes the smoothly time-varying assumption. *TGS-Lite* provides the same time complexity and true positive detection power as those of *TGS* at a significantly lower memory requirement, that grows linearly to the number of genes. Similarly, *TGS-Lite+* offers the superior time complexity and reconstruction power of *TGS+* with a linear memory requirement.

To summarise, the main contribution of this paper is three-fold:

- **Flexibility:** It provides a data-driven framework without imposing the smoothly time-varying assumption. In this framework, the time-varying GRN structures are reconstructed independently of each other. Thus, the framework is compatible with any time-series gene expression dataset, regardless of whether the true GRNs follow the smoothly time-varying assumption or not.
- **Time-efficiency:** The proposed framework offers state-of-the-art time complexity.
- **Memory-efficiency:** The memory requirement of the proposed framework grows linearly with the number of genes in the given dataset.

## 2 Problem Formulation

A time-varying GRN reconstruction algorithm takes a dataset 𝒟 as input and returns a sequence of time-varying GRNs 𝒢 as output (Figure 1). Dataset 𝒟 consists of *S* number of time series, denoted by 𝒮 = {*s*_1_*,…, s*_*S*_}. Each time series is comprised of the expression levels of *V* number of genes 𝒱 = {*v*_1_*,…, v*_*V*_} at *T* consecutive time points 𝒯 = {*t*_1_*,…, t*_*T*_}. 𝒟_(𝒳;𝒴;*Ƶ*)_ denotes the expression levels of genes 𝒳 at time points 𝒴 in time series *Ƶ*. 𝒟_(𝒳;𝒴;*Ƶ*)_ ⊆ 𝒟 since 𝒳 ⊆ 𝒱, 𝒴 ⊆ 𝒯, *Ƶ* ⊆ 𝒮. Dataset 𝒟 is also assumed to be complete i.e. there are no missing values.

**Fig. 1.**
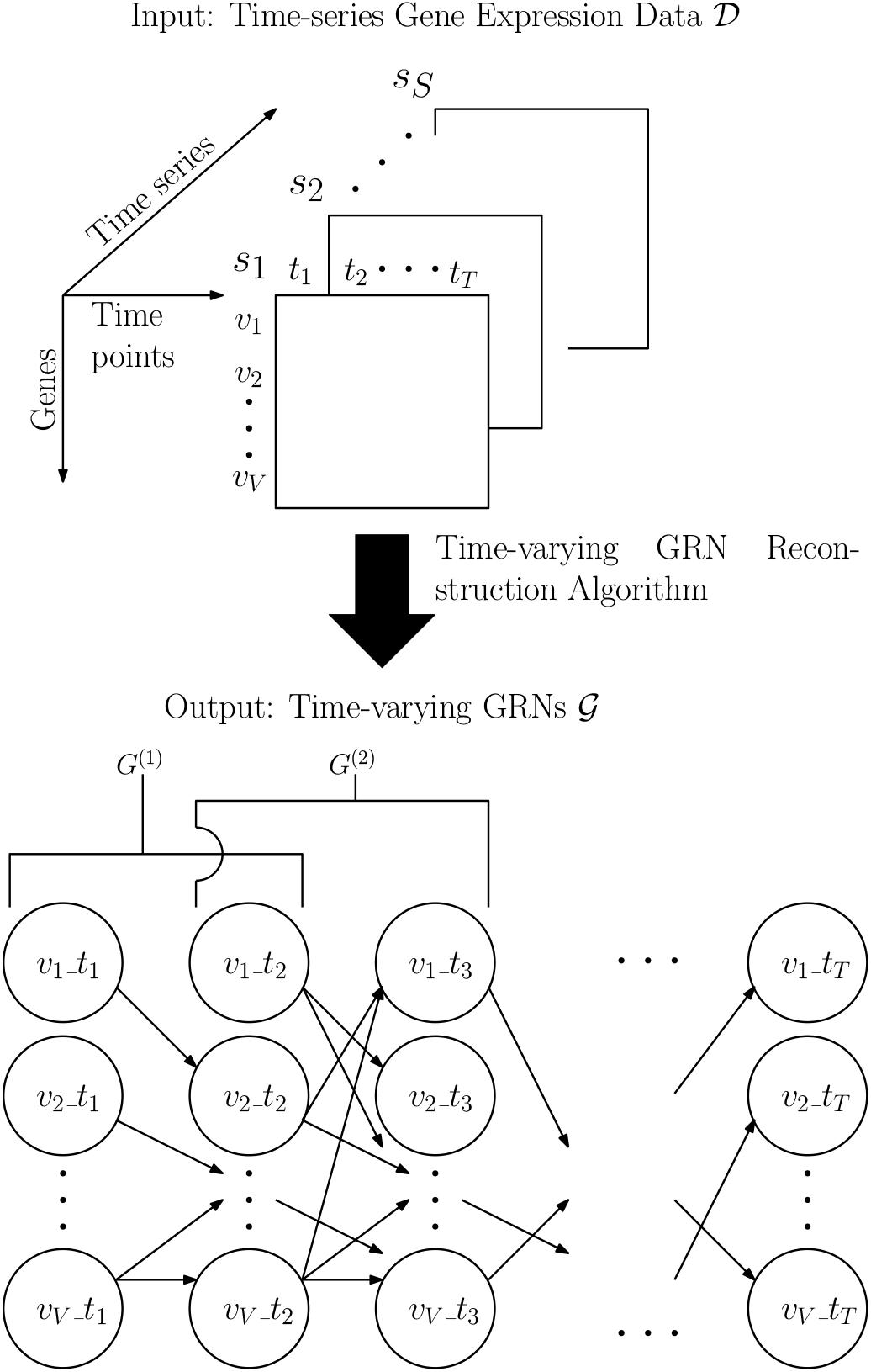
The Workflow of a Time-varying GRN Reconstruction Algorithm. The algorithm takes a time series gene expression data 𝒟 as input. The data consists of *S* number of time series. Each time series contains measured expressions of *V* number of genes across *T* number of time points. In return, the algorithm outputs time-varying GRNs (*G*^(1)^*,…, G*^(*T−*1)^) = 𝒢, which is a sequence of directed unweighted networks. Here, *G*^(*p*)^ (∈ 𝒢) represents the gene regulatory events occurred during the time interval between time points *t*_p_ and *t*_(*p*+1)_. It consists of (2 *× V*) nodes {*v*_*i*__*t*_*q*_: *v*_*i*_ ∈ 𝒱, *t*_*q*_ ∈ {*t*_*p*_, *t*_(*p*+1)_}}. There exists a directed unweighted edge (*v*_*i*__*t*_*p*_, *v*_*j*__*t*_(*p*+1)_ if and only if *v*_*i*_ regulates *v*_*j*_ during time interval (*t*_*p*_, *t*_(*p*+1)_). For instance, *G*^(1)^ represents the regulatory events that occurred between time points *t*_1_ and *t*_2_. One such event is the regulatory effect of *v*_1__*t*_1_ on *v*_2__*t*_2_ as represented by a directed edge.

Given dataset 𝒟, the objective is to reconstruct a temporally ordered sequence of GRNs 𝒢 = (*G*^(1)^;…;*G*^(*T*−1)^). Each *G*^(*p*)^ (∈ 𝒢) is a time interval specific GRN; it represents the gene regulatory events occurred during the time interval between time points *t*_*p*_ and *t*_(*p*+1)_. Structurally, *G*^(*p*)^ is a directed unweighted network with (2 × *V*) nodes: {*v*_i__*t*_*q*_: *v*_*i*_ ∈ 𝒱, *t*_*q*_ ∈ {*t*_*p*_, *t*_(*p*+1)_}}. Each node *v*_*i*__*t*_*q*_ is a distinct random variable, which represents the expression level of gene *v*_*i*_ at time point *t*_*q*_; hence, data points 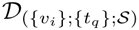 are considered to be *S* instances of random variable *v*_*i*__*t*_*q*_. It is assumed that the underlying gene regulation process is first order Markovian [5] i.e. node *v*_*i*__*t*_*q*_ can have regulatory effects only on the nodes at the immediately next time point *t*_(*q*+1)_, if any. Thus, there exists a directed edge (*v*_*i*__*t*_*q*_; *v*_*j*__*t*_(*q*+1)_) in 𝒢 if and only if *v*_*i*__*t*_*q*_ has a regulatory effect on *v*_*j*__*t*_(*q*+1)_, implying that the expression level of gene *v*_*i*_ at time point *t*_*q*_ plays a regulatory role on that of gene *v*_*j*_ at time point *t*_(*q*+1)_.

## 3 Existing Algorithms

There exists a set of algorithms that solve the problem partially, e.g., *Bene* [16], *GENIE3* [17], *NARROMI* [18], *LBN* [19]. They can reconstruct a time-invariant ‘summary’ GRN over node-set 𝒱. A directed edge (*v*_*i*_, *v*_*j*_) signifies that gene *v*_*i*_ is likely to have a regulatory effect on gene *v*_*j*_. However, it does not help in identifying the time interval(s) when that regulatory effect has taken place. On the other hand, Friedman et al. reconstruct one GRN for every time interval by modelling 𝒢 as a Dynamic Bayesian Network (DBN) [5]. However, DBN’s time-homogeneous nature requires each gene to have the same regulators in all time intervals.

To overcome time-homogeneity, is 𝒢 modelled as a time-inhomogeneous DBN by a group of algorithms ([3], [6], [7], [8], [9], [10], [13]). They assume that the underlying gene regulation process is a multiple-change-point process with the set of change points 𝔗 ⊆ 𝒯. The duration between two consecutive change points is called a time segment. Algorithms *NsDbn* [6] and *NsCdbn* [7] reconstruct a unique GRN for each time segment. A more flexible algorithm *ARTIVA* reconstructs a unique GRN for each time interval [8]. It assumes that every gene has unique change points. Thus, GRN structures of two consecutive time intervals vary if they belong to different time segments of at least one gene.

*ARTIVA*’s flexibility fetches two major criticisms. First, Grzegorczyk et al. suggest that having unique change points for every gene might be too flexible [3]. Instead, genes with similar expression patterns can be grouped into a cluster and unique change points can be assigned to that cluster. Based on this paradigm, Grzegorczyk et al. develop the *cpBGe* algorithm. However, *cpBGe* models 𝒢 as a directed weighted network whose edge weights are time-varying but the structure is time-invariant.

The next criticism comes from Dondelinger et al. [9], who become concerned of *ARTIVA*’s high computational cost since *ARTIVA* divides the reconstruction problem into a large number of atomic problems of learning the regulators of every gene during every time segment specific to that gene. Moreover, they argue that *ARTIVA* is statistically vulnerable to overfitting when *S ≪ V* since each atomic problem is solved only with the corresponding time segment’s data. To avoid these issues, Dondelinger et al. propose an alternative framework where all atomic problems are solved jointly through ‘information sharing’ or ‘coupling’. The underlying assumption, known as the ‘smoothly time-varying assumption’, is that every time-interval specific GRN structure shares more common edges with its temporally adjacent GRN structures than with the distal ones. All atomic problems are solved jointly to ensure that the reconstructed GRN structures honour the assumption. The framework is further divided into two categories – soft coupling and hard coupling – based on the strength of coupling i.e. the expected amount of similarities between the GRN structures. To realize this framework, Dondelinger et al. first introduce a baseline algorithm without coupling, namely *TVDBN-0*, which is similar to *ARTIVA* except in its internal sampling strategy. Then, coupling strategies are added to the baseline algorithm to develop two hard coupling algorithms - *TVDBN-bino-hard* and *TVDBN-exp-hard*, and two soft coupling algorithms – *TVDBN-bino-soft* and *TVDBN-exp-soft*; the terms ‘bino’ and ‘exp’ represent that the corresponding algorithm assumes the expression levels of every gene to follow a binomial or an exponential distribution, respectively. Based on the same smoothly time-varying assumption, one more algorithm (henceforth, *MAP-TV*) is proposed by Chan et al. [10], which is later extended by Zhang et al. [13].

A comparative study between *ARTIVA* and the latter algorithms is conducted by Pyne et al. [14]. When compared against three benchmark datasets, they observe that *ARTIVA* most consistently provides the most correct models i.e. the highest F1-scores. Two potential reasons behind this observation are: (1) the smoothly time-varying assumption does not hold for the given datasets and (2) each of the datasets possesses a sufficient number of time series which nullifies the *S ≪ V* condition. On the other hand, it is also observed that *ARTIVA* consumes the largest amounts of time. Inspired by these observations, Pyne et al. develop a time-efficient framework which is also as flexible as *ARTIVA*. This framework considers every time point as a change point for every gene. Thus, solving the reconstruction problem boils down to solving a large number of atomic problems of identifying the regulators of every gene during every time interval. Instead of attempting to reduce the number of atomic problems, Pyne et al. propose to reduce the amount of time consumed by each atomic problem. Since the time complexity of each atomic problem grows exponentially with the number of candidate regulators, an information theoretic strategy is applied to significantly shorten that number. Two algorithms – *TGS* and *TGS+* – are conceived based on this framework. *TGS* most consistently outperforms *ARTIVA* and other algorithms in true positive detection power as well as in computational speed. However, *ARTIVA* retains its superiority in false positive rejection power. *TGS+*, on the other hand, consistently provides a true positive detection power comparable to that of *TGS* and a false positive rejection power comparable to that of *ARTIVA*. As a result, *TGS+* replaces *ARTIVA* in most consistently achieving the highest F1-scores. Moreover, it replaces *TGS* for being the fastest. Nevertheless, the flexibility and time-efficiency of *TGS* and *TGS+* come at the cost of their memory requirements, which grow exponentially to the number of genes. For these reasons, developing a flexible, time-efficient as well as memory-efficient framework can be considered as a timely contribution. (Please find a summary of the discussed algorithms at Table 1.)

**TABLE 1.** A Summary of the Existing Algorithms, discussed in Section 3.

| Algorithm(s) | Summary |
| --- | --- |
| *Bene* [16], *GENIE3* [17], *NARROMI* [18], Chang et al. [11], *LBN* [19], *cpBGe* [3] | Learn a single time-invariant GRN structure (the ‘summary GRN’). Hence, can not identify the time interval during which a regulatory event has occurred. |
| Friedman et al. [5] | Learns time-varying GRNs. However, requires each gene to have the same regulators in all time intervals. |
| *NsDbn* [6], *NsCdbn* [7], *ARTIVA* [8], Xiong et al. [4], *TVDBN-0* [9] | Allow each gene to have different regulators at different time intervals. Nonetheless, substantially slow. |
| {*TVDBN-bino-hard*, *TVDBN-bino-soft*, *TVDBN-exp-hard*, *TVDBN-exp-soft*} [9], *MAP-TV* [10], Zhang et al. [13] | Relatively faster. Nevertheless, impose the structural constraint that each GRN shares more common edges with its temporally adjacent GRNs than the distal ones. This constraint is known as the ‘smoothly time-varying assumption’. |
| {*TGS*, *TGS+*} [14] | The fastest. Do not require the smoothly time-varying assumption. However, exponential memory requirements. |

## 4 Methods

In this section, the objective is to develop novel algorithms that have equivalent learning power and speed to *TGS* and *TGS+* but significantly less memory requirements. For that purpose, first, investigations are conducted to identify the origin of the exponential memory requirements of *TGS* and *TGS+*. Then novel algorithms are designed to overcome this issue.

### 4.1 Investigations into the Origin of the Issue

An analytical study of the *TGS* algorithm with the max fan-in^1^ restriction (‘Algorithm 4’ in Pyne et al. [14]) reveals the origin of its high memory requirement. *TGS* has two distinct steps. In the first step, for each node *v*_*j*__*t*_(*p*+1)_ in 𝒢, it generates a candidate regulator set, denoted by 𝒱_(*j*;(*p*+1))_. In the second step, it chooses the best set of regulators for *v*_*j*__*t*_(*p*+1)_ from 𝒱_(*j*;(*p*+1))_. The issue originates from the second step (hereafter, the Bene step) where *TGS* employs a Bayesian Network (BN) structure learning algorithm, namely *Bene*, to choose the best set of regulators. *Bene* does so by calculating BIC scores [20] of all subsets of 𝒱_(*j*;(*p*+1))_ and choosing the subset with the highest BIC score. For that purpose, the Bene step requires 2^|𝒱_(*j*;(*p*+1))_|^ scores and 2^|𝒱_(*j*;(*p*+1))_|^ subsets to be held in memory (Section 4.1 of the supplementary document). It results in an exponential memory requirement w.r.t. the max fan-in parameter *M*_*f*_, since |𝒱_(*j*;(*p*+1))_| = 𝒪(*M*_*f*_).

### 4.2 A Novel Idea for Resolving the Issue

A more memory-efficient algorithm (Algorithm 1) is designed to replace the Bene step. It requires two pointers (curr.set and best.set), only two scores (curr.score and best.score) and 2^|𝒱_(*j*;(*p*+1))_|^ subsets to be held in memory (Figure 2A). Nevertheless, the number of subsets in memory remains exponential w.r.t. *M*_*f*_.

**Fig. 2.**
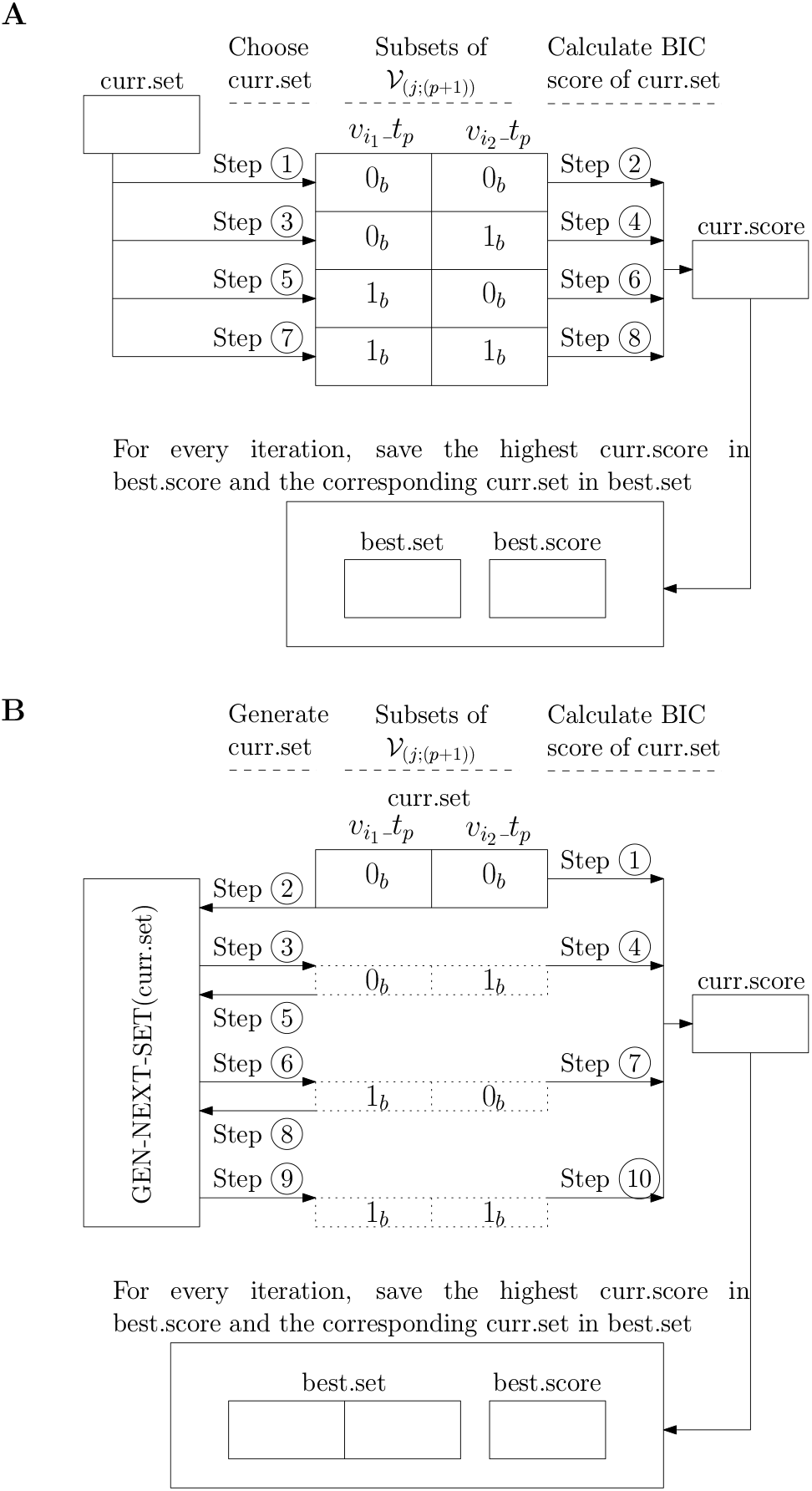
Memory Footprints of Algorithm 1 and Algorithm 3: a comparison. For illustration, it is assumed that 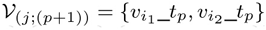. Therefore, its subsets are 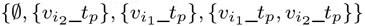. Each subset is represented as a binary string of Boolean TRUE (1_*b*_) and FALSE (0_*b*_) values, indicating whether a gene is present or not in that subset, respectively. **A)** Memory footprint of algorithm 1: the algorithm requires two pointers (curr.set and best.set), two scores (curr.score and best.score) and 2^|𝒱_(*j*;(*p*+1))_|^ = 2^2^ = 4 subsets to be held in memory. **B)** Memory footprint of algorithm 3: it requires a subset-generation script (GEN-NEXT-SET), two scores (curr.score and best.score) and only two subsets (curr.set and best.set) to be held in memory. Here, curr.set holds a subset whereas it holds a pointer to a subset in case of algorithm 1. In algorithm 3, the value of curr.set is updated to the next subset by the GEN-NEXT-SET script through in-place replacement, saving memory. Both the algorithms also require data 𝒟* (not shown in figure) to be stored in memory during the calculation of BIC scores.

One naive way to reduce the number of subsets in memory is to keep only the subsets pointed by curr.set and best.set in memory and move the rest of the subsets to the secondary storage (hereafter, disk). However, that strategy will increase runtime significantly due to costly disk I/Os.

Keeping all the subsets in memory or performing disk I/Os – both can be avoided if every subset can be generated in real-time only when its BIC score needs to be calculated. From Figure 2A, it is observed that every non-empty subset can be generated by adding 1_*b*_ to the Least Significant Bit of its previous subset (Algorithm 2). Using this strategy, a novel algorithm, namely *Find-best-set-Lite* (Algorithm 3), is designed to find the highest scoring subset. It requires a subset-generation script, only two scores (best.score and curr.score) as well as only two subsets (curr.set and best.set) to be held in memory (Figure 2B), resulting in a linear memory requirement w.r.t. *M*_*f*_.

### 4.3 Design of Novel Algorithms

#### TGS-Lite

A novel algorithm, namely ‘an algorithm for reconstructing Time-varying Gene regulatory networks with Shortlisted candidate regulators - which is Light on memory’ or *TGS-Lite* (Algorithm 4), is conceived by replacing the Bene step in *TGS* with *Find-best-set-Lite*. Thus, *TGS-Lite* is able to achieve a significantly lower memory footprint compared to that of *TGS* while reconstructing the same time-varying GRNs. At the same time, *TGS-Lite*’s time complexity (Equation 4.8 of the supplementary document) remains same as that of *TGS* with the max fan-in restriction (Equation 4.13 of the supplementary document) i.e. *o* (*V*^2^ lg *V*).

#### TGS-Lite+

Since *TGS-Lite* reconstructs the same GRNs as *TGS*, it suffers from the same issue of high false positives. A variant of *TGS-Lite*, namely ‘TGS-Lite Plus’ (*TGS-Lite+*), is designed to mitigate this issue (Algorithm 5). The only difference in this new variant is the addition of the ARACNE step (Algorithm 5 lines 7 - 9), which is shown to significantly reduce false positives at a reasonable reduction in true positives [14]. Although the addition of the ARACNE step increases the time complexity (Section 4.5 of the supplementary document), it helps to produce a shorter list of candidate regulators for each gene [14], thus saving time during the final selection of regulators.

## 5 Results

In this section, two sets of experimental results are presented for *TGS-Lite* and *TGS-Lite+*: one with three in-silico bench-mark datasets and the other with a real microarray dataset.

### Algorithm 1 Find-best-set

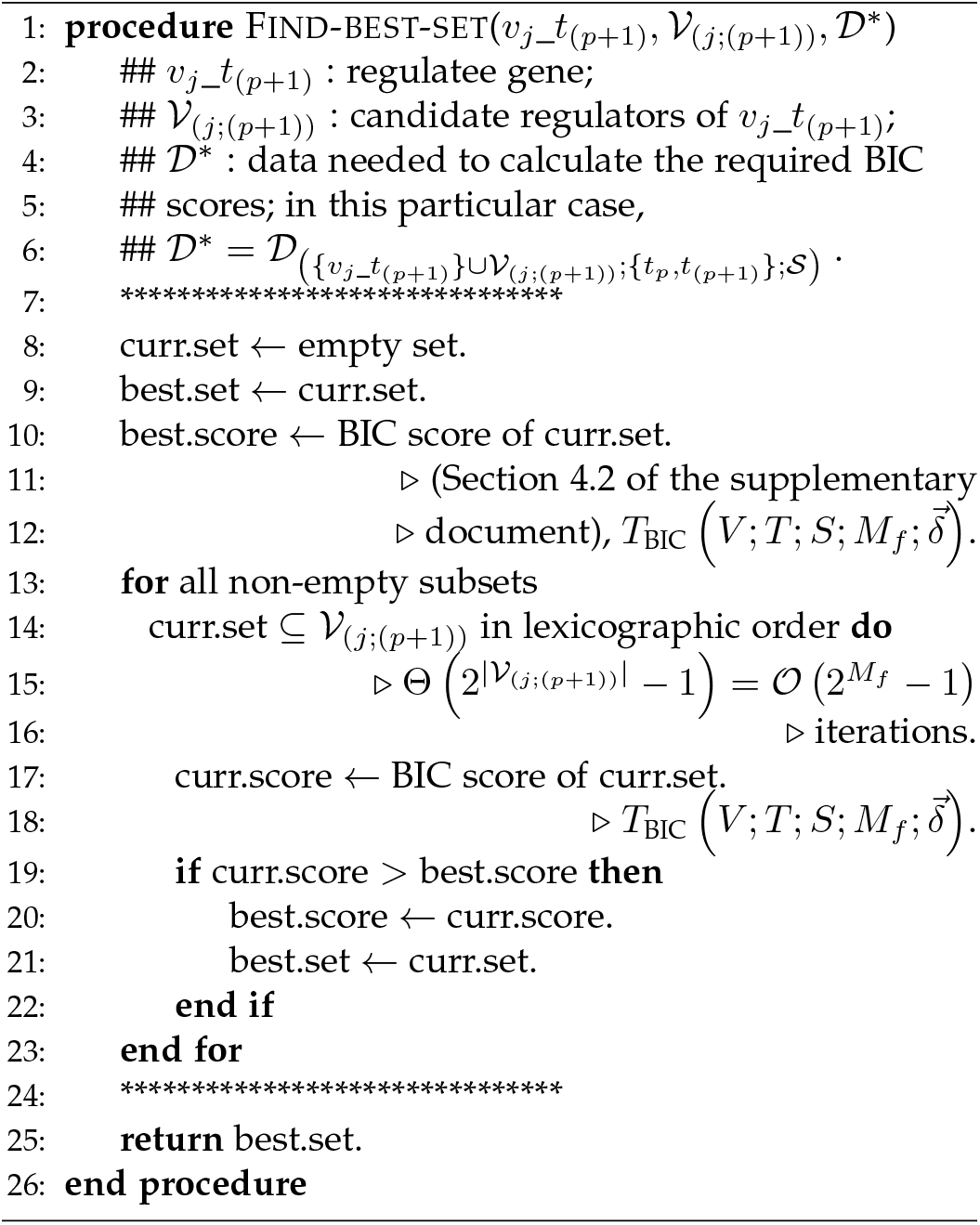

### 5.1 Results with In-silico Benchmark Datasets

The in-silico benchmark datasets used for this study are known as Ds10n, Ds50n and Ds100n [14]. They are originally published through DREAM3 In Silico Network Challenge ([21], [22], [23], [24]) for systematically comparing GRN reconstruction algorithms. The challenge also published the ‘true’ GRNs that generated the datasets. Although the datasets are longitudinal, the true GRNs are time-invariant in nature. A brief description of these datasets and corresponding true GRNs are given in Table 2. Their detailed description can be found in Pyne et al. [14]. Since the true GRNs are time-invariant and the predicted GRNs are time-varying, the predicted GRNs are ‘rolled’ into time-invariant GRNs to determine their correctness. The rolling strategy is a union operation over edges i.e. a directed edge from gene *v*_*i*_ to gene *v*_*j*_ exists in the rolled GRN if and only if the edge exists in at least one of the time-varying GRNs. Since {*TGS-Lite, TGS-Lite+, TGS, TGS+*} require discretised data, the 2L.wt algorithm [14] is used for data discretisation. The metrics used for evaluating the correctness of the predicted GRNs are described in Section 4.8 of the supplementary document.

**TABLE 2.** A Summary of the chosen DREAM3 Datasets. V = number of genes. T = number of time points. S = number of time series.

| Dataset | V | T | S | No. of True Edges |
| --- | --- | --- | --- | --- |
| Ds10n | 10 | 21 | 4 | 10 |
| Ds50n | 50 | 21 | 23 | 77 |
| Ds100n | 100 | 21 | 46 | 166 |

#### 5.1.1 Preliminary Study against a Random Classifier

Before moving towards a comprehensive analysis, a preliminary study is conducted to check whether the results of *TGS-Lite* and *TGS-Lite+* are better than random or not.

##### Algorithm 2 Gen-next-set

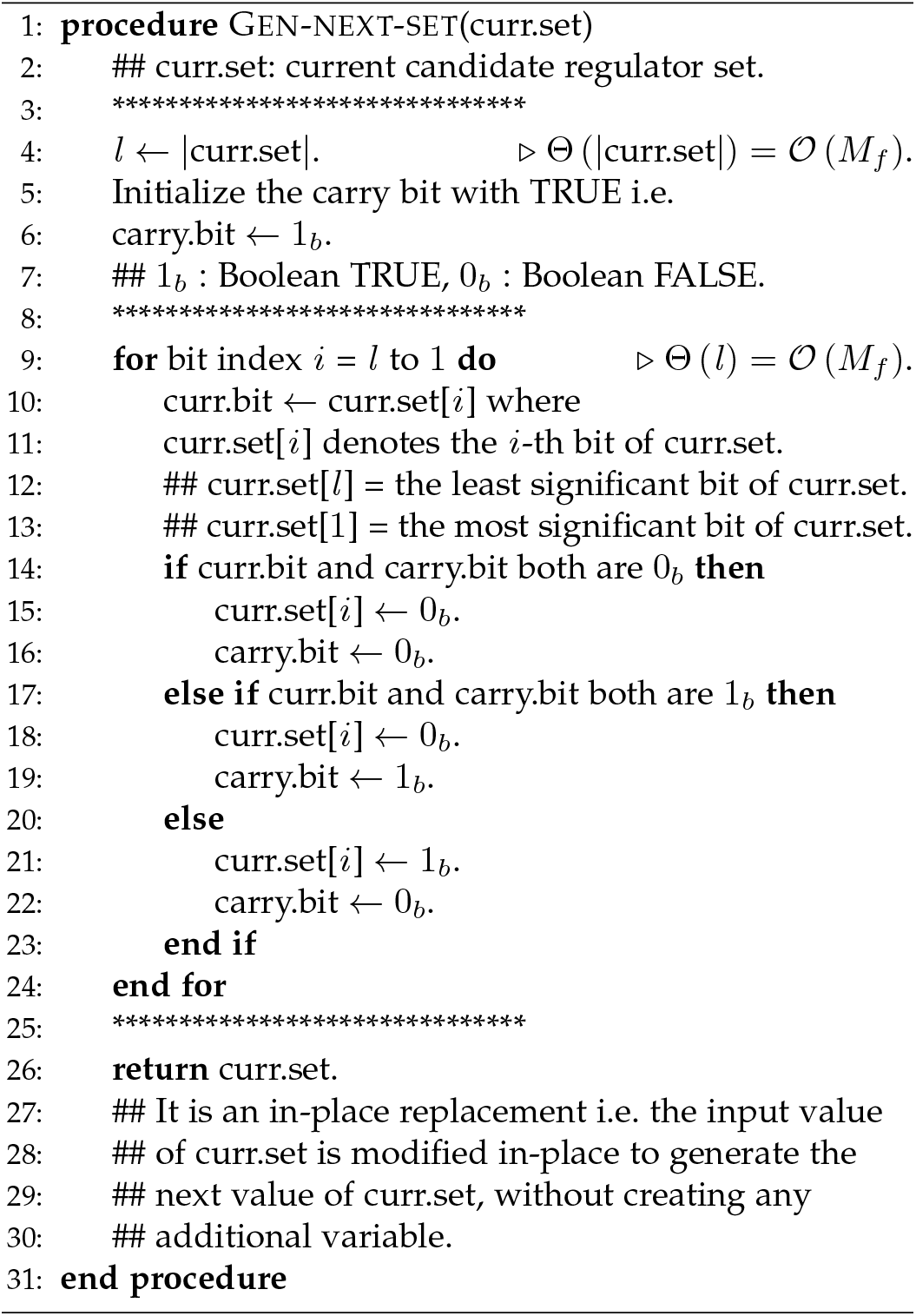

For that purpose, TPR-vs-FPR chart of *TGS-Lite* and *TGS-Lite+* is plotted (Figure 3). These plots remain above the random classifier’s line, signifying that the results are better than random. Here, the random classifier line represents the results of a classifier that randomly decides whether an edge should be present or absent in the predicted network. Such random classification tends to result in a line represented by the equation TPR = FPR.

**Fig. 3.**
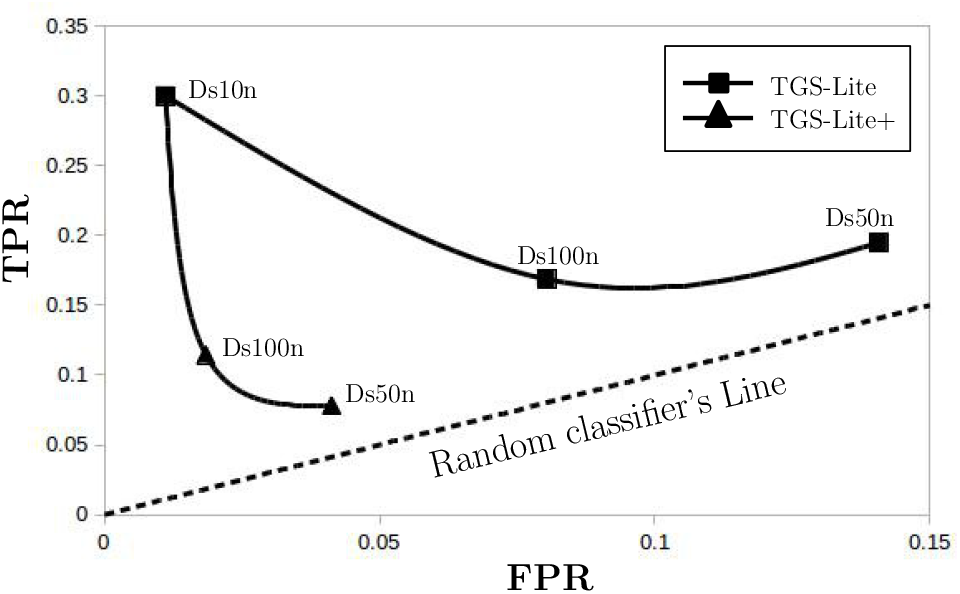
The TPR-vs-FPR Plots of *TGS-Lite*, *TGS-Lite+* and a random classifier. Here, TPR = True Positive Rate, FPR = False Positive Rate. The results of *TGS-Lite* for three distinct DREAM3 datasets are represented as three black squares. These squares are connected (interpolated) with a smooth line (Software used: LibreOffice Calc Version 5.1.6.2; Line type = Cubic spline, Resolution = 20; OS: Ubuntu 16.04.5 LTS). On the other hand, the results of *TGS-Lite+* for three distinct DREAM3 datasets are represented as three black triangles (the triangle for Ds10n overlaps with the square for Ds10n and hence not visible). These triangles are also connected with a smooth line.

#### 5.1.2 Comparative Study against Alternative Algorithms

##### Experimental Settings

Following the success of *TGS-Lite* and *TGS-Lite+* in the preliminary test, a comparative study is conducted against a set of alternative algorithms, which are {*TGS*, *TGS+*, *ARTIVA*, *TVDBN-0*, *TVDBN-bino-hard*, *TVDBN-bino-soft*} (Figure 4). The results of these alternative algorithms are reproduced from Pyne et al. [14] since the same hardware and Operating System are used. The only difference is that those algorithms are interpreted in R programming language [25] version 3.3.2 and {*TGS-Lite*, *TGS-Lite+*} are interpreted in R version 3.5.1. For each of {*TGS-Lite*, *TGS-Lite+*, *TGS*, *TGS+*}, the naming convention used are as follows: no extension = serial execution with *M*_*f*_ set to 14; ‘mf[X]’ = *M*_*f*_ set to ‘X’, e.g., *TGS.mf24*; ‘p[X]’ = parallel execution with number of cores set to ‘X’, e.g., *TGS-Lite.p10*.

###### Algorithm 3 Find-best-set-Lite

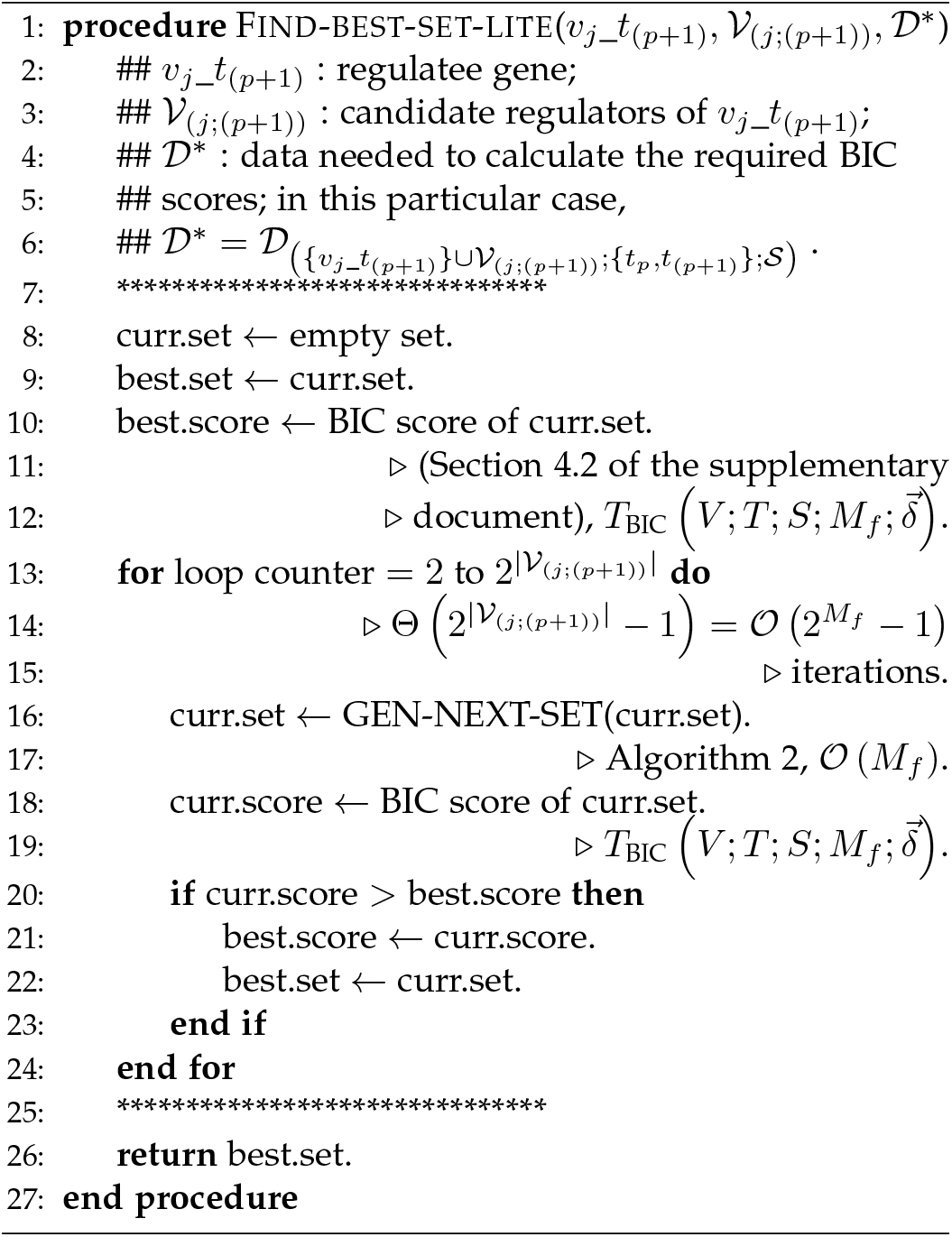

###### Algorithm 4 *TGS-Lite* with the Max Fan-in Restriction

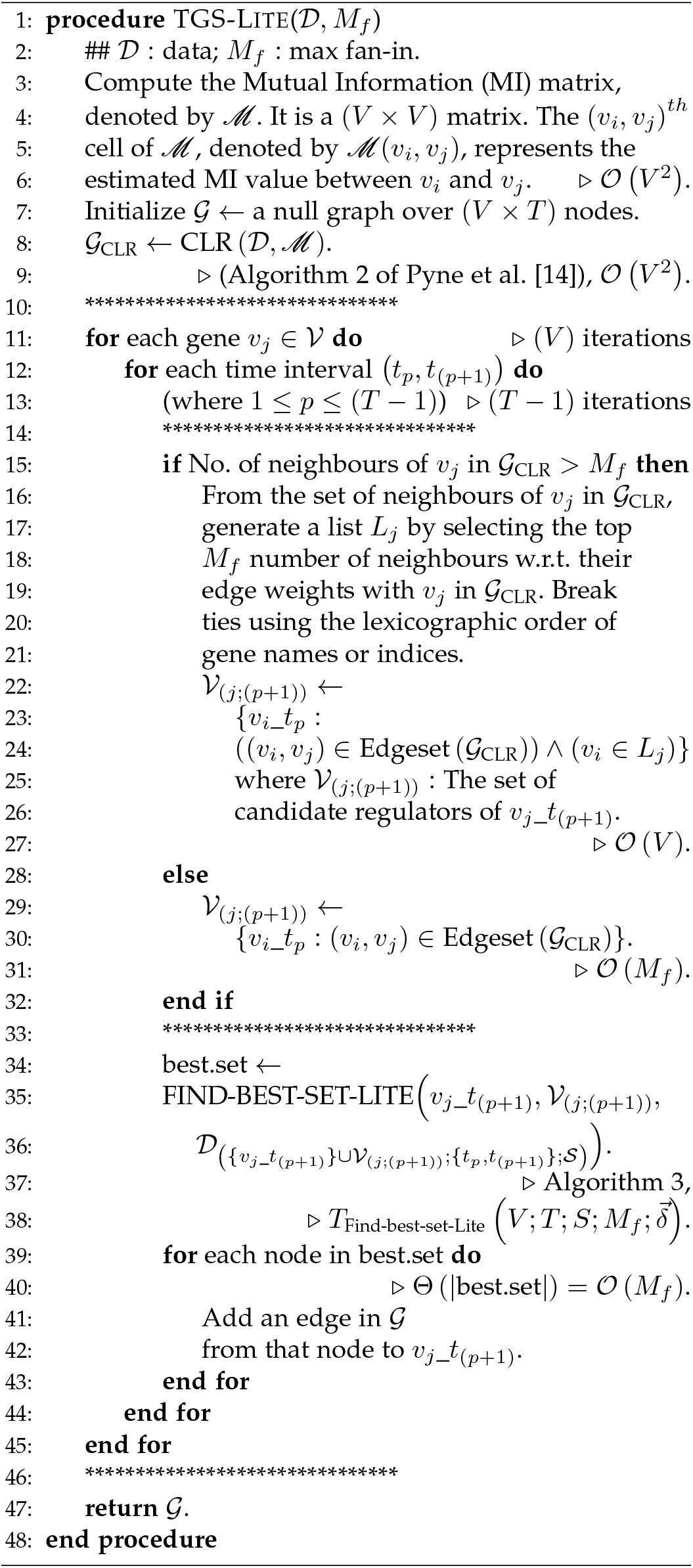

**Fig. 4.**
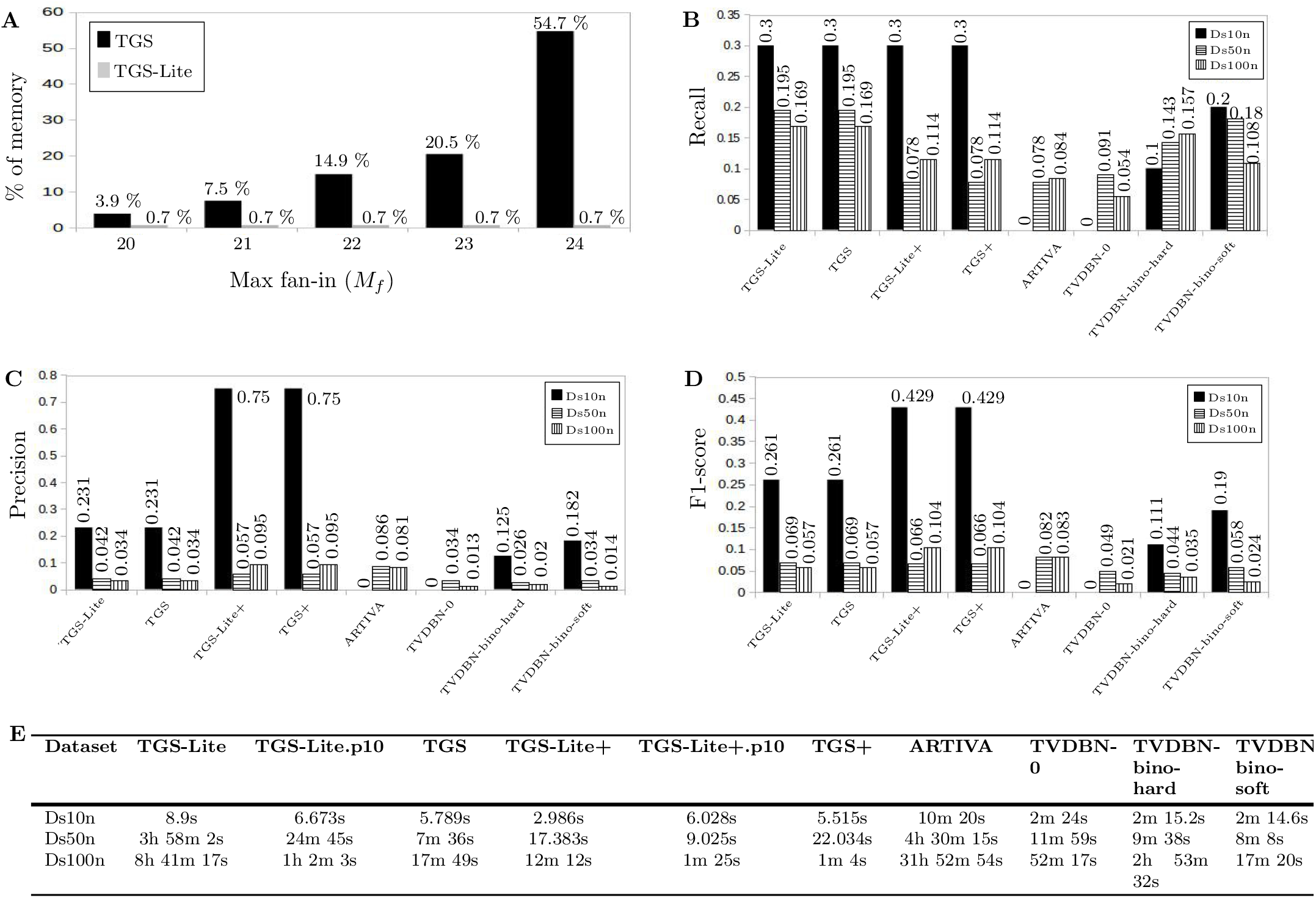
Comparative Performances of the selected Algorithms on the DREAM3 datasets. **A)** The percentages of memory used by *TGS* and *TGS-Lite* for different max fan-in values. The percentages represent memory usage during the beginning of the Bene step (see Section 4.11 of the supplementary document for more details). Dataset in use is Ds100n. **B)** Recall, **C)** precision, **D)** F1-score and **E)** runtime of the selected algorithms are shown. *Recall = TP / (TP + FN); Precision = TP / (TP + FP); F1-score = (2 × Recall × Precision) / (Recall + Precision); here, TP or True Positive = number of true edges correctly predicted by the concerned algorithm; FN or False Negative = number of true edges that are not predicted; FP = number of predicted edges that are not true*.

##### Lighter Memory Footprint

Figure 4A demonstrates a significant reduction in memory usage in case of *TGS-Lite* compared to *TGS*. For *M*_*f*_ = 24, TGS occupies 54.7% of the main memory (~32 GB) during the beginning of the Bene step. Its memory usage rises to almost 100% as the time goes on and finally reaches a ‘thrashing’ state [26]. At that point, the *TGS* process is terminated. In comparison, *TGS-Lite* occupies a negligible amount of memory (0.7%) and completes execution without any issue. Moreover, it indicates the possibility of parallel executions with at most (100/0.7) ≃ 142 cores, significantly reducing runtime as well. On the other hand, the heavy memory footprint of *TGS* prevents it from taking advantage of such multicore parallelisation schemes.

##### No Loss in the Learning Power

The memory efficiency of *TGS-Lite* does not come at the cost of its learning power. It provides the same recall, precision and F1-score as those of *TGS*. Together they provide the highest recalls for all three datasets (Figure 4B). In case of precision, only *TGS-Lite+* and *TGS+* are able to outperform them for all datasets (Figure 4C). Owing to such high precisions, coupled with competitive recalls, *TGS-Lite+* and *TGS+* jointly obtain the highest F1-scores for two out of three datasets (Figure 4D). Only for Ds50n, *ARTIVA* provides the highest F1-score, outperforming {TGS-Lite+, TGS+} by a margin of 0.016; however for the other two datasets, the latter algorithms supersede *ARTIVA* by larger margins (0.429 and 0.021). Interestingly, only for Ds10n, *TGS-Lite+* retains the recall obtained by *TGS-Lite* (Figure 4B). The reason is that the lower the number of feed-forward edges in the true network, the lower the chances of the ARACNE step missing true edges [14]. Since, the true networks of Ds50n and Ds100n have large numbers of feed-forward edges (≃39%), the ARACNE step misses a large number of true edges. However, for Ds10n, whose true network has 10% feed-forward edges, it misses only one true edge. The ARACNE step compensates this small loss by helping *TGS-Lite+* in capturing a true edge that *TGS-Lite* misses. Thus, for Ds10n, *TGS-Lite+* predicts as many true edges as *TGS-Lite’s* (see Section 4.10 of the supplementary document for more details).

##### Multicore Parallelisation

Although *TGS-Lite* shares the same time complexity with *TGS*, it encounters larger runtime than those of *TGS* (Figure 4E). The potential reason being the difference in their implementations. While the Find-best-set-Lite step in *TGS-Lite* is implemented with R (except the function for BIC score computation which is in C), its counterpart in *TGS*, the Bene step, is implemented with C, which is expected to be significantly faster. To alleviate this issue, another R implementation of *TGS-Lite* is prepared with multicore parallelisation. The 10-core parallelised execution of *TGS-Lite* i.e. *TGS-Lite.p10* reduces the runtime by a factor of eight for larger datasets: Ds50n and Ds100n (Figure 4E). An implementation of *TGS-Lite* with C is planned for future which is likely to be even faster.

###### Algorithm 5 *TGS-Lite+* with the Max Fan-in Restriction

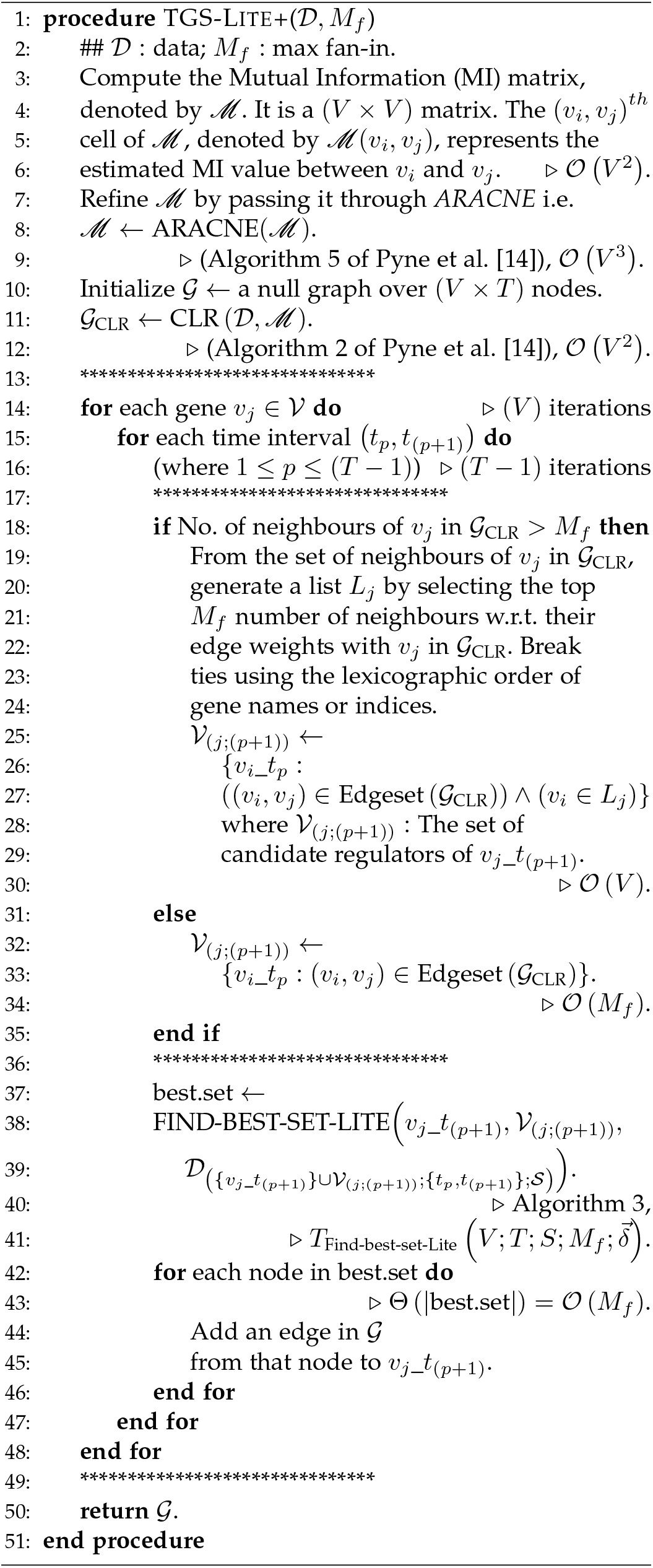

*TGS-Lite+* finds a right amount of balance between the memory efficiency and computational speed while providing the superior F1-score (Figure 4D). Its low runtime make it the second fastest algorithm, slightly slower than *TGS+* (Figure 4E) but without *TGS+*’s heavy memory footprint [14]. The 10-core parallelised execution of *TGS-Lite+* (TGS-Lite+.p10) decreases the runtime further (Figure 4E). However, there is an interesting exception: for Ds10n, the parallel execution takes more time than that of the serial execution. This is because the dataset is so small that the communication overhead with multiple cores outweighs the reduction in runtime due to parallelisation. For the larger two datasets, parallelisation saves time, making the runtime competitive to that of *TGS+*. Therefore, the parallel implementation of *TGS-Lite+* can be advantageous when there is a large number of genes (50+) while the serial implementation is sufficiently fast for smaller datasets.

##### Effect of Multicore Parallelisation

This paragraph studies the effect of the multicore parallelisation on runtime of *TGS-Lite* and *TGS-Lite+* in more depth. The dataset chosen for this study is Ds100n since it is the largest dataset. For both the algorithms, the runtime strictly decrease with the increase in the number of cores (Figure 5). The results with a single core represent the serial executions, which can be used as a baseline. Following that, the speed-up with three and seven cores could be of interest to users with quad-core and octa-core processors, respectively, since one core can be left out for monitoring purposes.

**Fig. 5.**
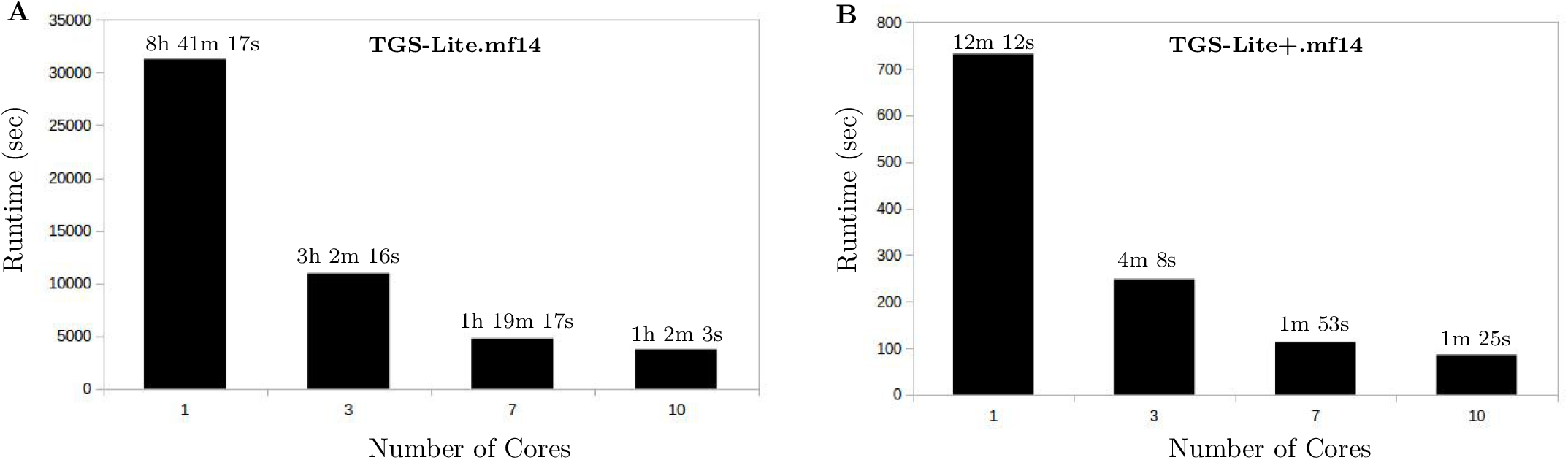
Effect of the Multicore Parallelisation on the Runtime of *TGS-Lite* and *TGS-Lite+*. Dataset Ds100n is used. Max fan-in is set to 14.

##### Effect of Max Fan-in

In this paragraph, the effect of the max fan-in parameter on the learning power and speed of the *TGS-Lite* and *TGS-Lite+* algorithms is studied. The objective of this study is to set max fan-in for *TGS-Lite* and *TGS-Lite+* to values higher than 14, which is the largest value with which *TGS* and *TGS+* is hitherto studied without any segmentation fault [14]. Dataset Ds100n is chosen for that purpose since it is the only dataset for which the maximum number of neighbours in _CLR_ of both *TGS-Lite* and *TGS-Lite+* exceeds 14 (Table 3). Therefore, the results (Figure 6) are obtained by varying max fan-in from 14 to *min*(84, 18) = 18. For *TGS-Lite*, the recall monotonically increases with the increase in max fan-in; however, the precision monotonically decreases, causing an upheaval in F1-scores. On the other hand, *TGS-Lite+* demonstrates a relatively robust performance, since its recall, precision and F1-score remain unchanged for the given range of max fan-in. The effect of max fan-in on runtime is found to be proportional for both the algorithms. This observation complies with their time complexities.

**TABLE 3.** Maximum Number of Neighbours of a Gene in 𝒢_CLR_ of the *TGS-Lite* and *TGS-Lite+* algorithms for a given dataset.

| Dataset | TGS-Lite | TGS-Lite+ |
| --- | --- | --- |
| Ds10n | 7 | 4 |
| Ds50n | 33 | 8 |
| Ds100n | 84 | 18 |

**Fig. 6.**
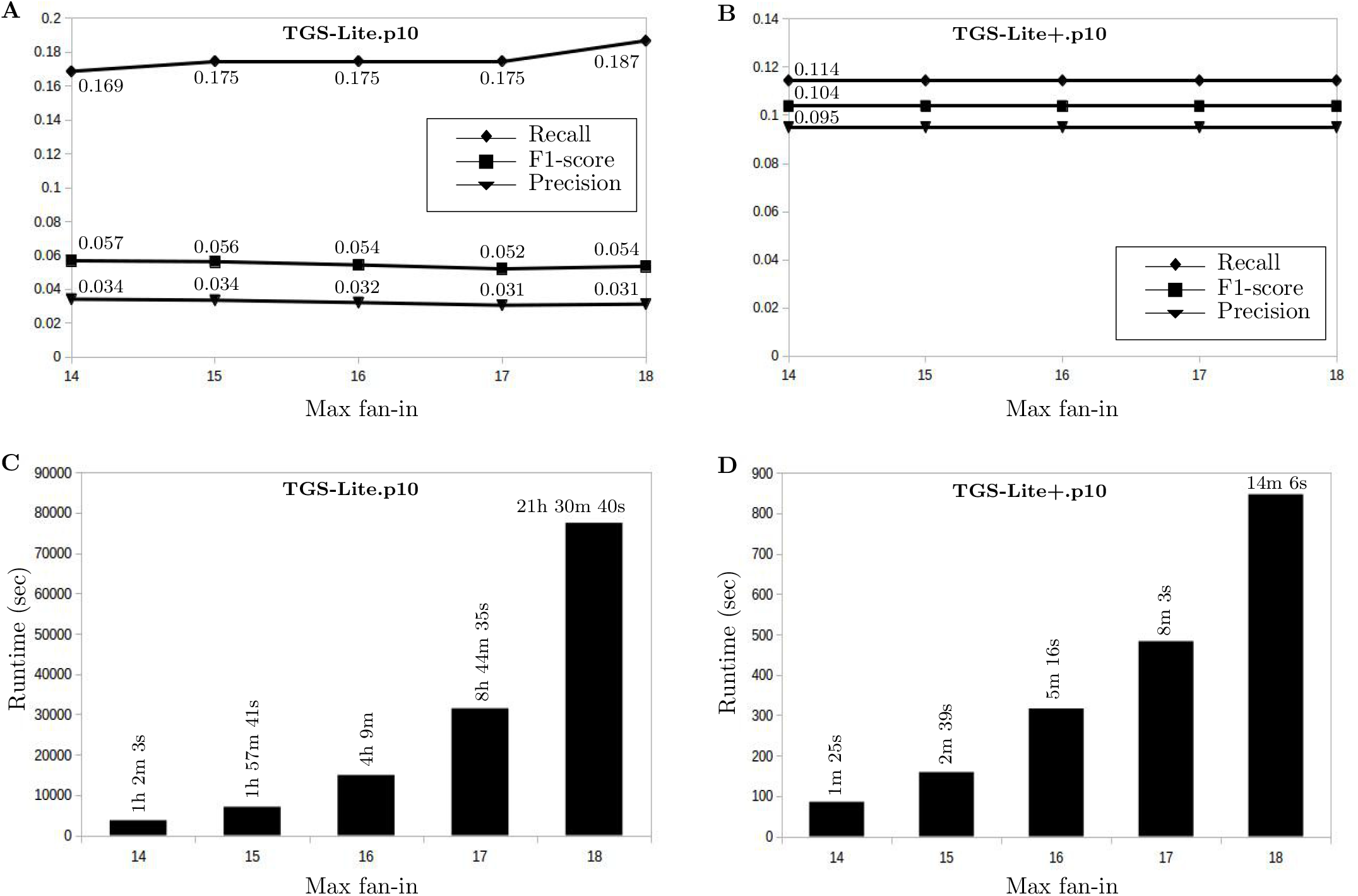
Effect of Max Fan-in on the Learning Power (**A, B**) and Speed (**C, D**) of the TGS-Lite and TGS-Lite+ algorithms. Dataset Ds100n is used and number of cores is set to 10 for multicore parallelisation.

##### A Comparison with the DREAM3 Winner

An additional comparative study is conducted against the performance of the winning team’s algorithm (hereafter, ‘*BTA*’) in DREAM3 In Silico Network Challenge [27]. Since, *BTA* reconstructs a summary GRN, it is compared against the rolled GRNs of {*TGS-Lite*, *TGS-Lite+*}. The comparison shows that *BTA* makes the highest true positive predictions while also conceding the highest false positives. On the contrary, *TGS-Lite+* provides the best balance between true and false positives; moreover, it achieves the lowest runtime. This study strengthens the superiority of *TGS-Lite+* (see Section 4.9 of the supplementary document for details).

### 5.2 Results with a Real Microarray Dataset

The real microarray dataset used in this study is known as DmLc3 [14]. It is originally produced by Arbeitman et al. [28]. DmLc3 contains gene expressions of the Drosophila melanogaster (Dm; fruit fly) life cycle. It is comprised of four sub-datasets corresponding to four Dm life cycle stages: DmLc3E (embryonic stage), DmLc3L (larval stage), DmLc3P (pupal stage) and DmLc3A (adulthood) (Table 4). Each subdataset contains the same 588 genes known to be involved in the developmental process of Dm according to their Gene Ontology (GO) annotations [29]. These sub-datasets are used to study the learning powers of *TGS* and *TGS+* in Pyne et al. [14].

**TABLE 4.** A Summary of the DmLc3 Sub-datasets. V = number of genes. T = number of time points. S = number of time series.

| Sub-dataset | V | T | S |
| --- | --- | --- | --- |
| DmLc3E | 588 | 6 | 5 |
| DmLc3L | 588 | 2 | 5 |
| DmLc3P | 588 | 3 | 6 |
| DmLc3A | 588 | 2 | 4 |

#### Evaluation Strategy

Since true GRNs are not known, the authors follow the same strategy as Pyne et al. [14] to evaluate the predicted GRNs. The strategy is based on a chosen subset of 25 genes that are known to play regulatory roles in Dm development. Hitherto experimentally verified knowledge about these genes is retrieved from TRANSFAC Public Database version 7.0 [30], which is claimed to be the gold standard in the area of transcriptional regulation [31]. The strategy is to find out whether these genes are predicted to have regulatory roles (at least one regulatee) in the known stages and whether they are predicted to regulate their known regulatees, if any.

#### Selection of Max Fan-in

Pyne et al. [14] restricts the max fan-in to 14 to avoid segmentation faults for *TGS* and *TGS+*. In this study, the authors set the max fan-in to 15 since *TGS-Lite* and *TGS-Lite+* do not require that restriction. The primary objective of this study is to find out whether *TGS-Lite.mf15* can identify more true edges than *TGS.mf14* does. The secondary objective is to assess whether *TGS-Lite+.mf15* can identify as many true edges as *TGS-Lite.mf15* while incurring less false positives. However, it must be noted that the prior knowledge only consists of a subset of true edges. Hence, it is not possible to mark false positive predictions. Therefore, the secondary objective is revised to assess whether *TGS-Lite+.mf15* can identify as many true edges as *TGS-Lite.mf15* while incurring a less number of potentially false positive edges. Some of the findings are discussed below (see Table 3.1 of the supplementary document for the complete set of findings).

#### Gene ‘Antp’

‘Antp’ is known to be essential for defining embryonal segment identity. In the embryonic stage, *TGS-Lite.mf15* identifies three regulatees of ‘Antp’ — ‘exu’, ‘opa’ and ‘aft’ — in addition to those identified by TGS.mf14. Identification of these additional regulatees is a potential true positive prediction since the regulatees are co-localized with ‘Antp’ in nucleus according to their Gene Ontology (GO) annotations. Especially, ‘opa’ is highly likely to be a regulatee of ‘Antp’ since ‘opa’ acts as a pair-rule gene (a gene involved in the development of segmented embryos) during early embryogenesis; see Sections ‘Gene Snapshot’ and ‘GO Summary Ribbons’ of ‘Antp’ (http://flybase.org/reports/FBgn0260642), ‘exu’ (http://flybase.org/reports/FBgn0000615), ‘opa’ (http://flybase.org/reports/FBgn0003002) and ‘aft’ (http://flybase.org/reports/FBgn0026309). *TGS-Lite+.mf15* also agrees with this prediction, improving confidence in ‘opa’ to be a direct regulatee of ‘Antp’. The experimental validation of this prediction can be considered as a novel opportunity.

#### Gene ‘eve’

‘eve’ is known to contribute to the development of the central nervous system. The regulatees of ‘eve’ predicted by *TGS-Lite.mf15* agree with those of *TGS.mf14* with one exception: ‘capu’ is replaced with ‘mmy’ as a regulatee of ‘eve’ in the embryonic stage. The ‘eve’ to ‘mmy’ edge is potentially a true positive prediction since ‘mmy’ is known to regulate axon guidance, which is an essential part of neural development. Hence, it is highly likely that there is a crosstalk between ‘eve’ and ‘mmy’; see Section ‘Gene Snapshot’ of ‘eve’ (http://flybase.org/reports/FBgn0000606) and ‘mmy’ (http://flybase.org/reports/FBgn0259749). On the other hand, the rejection of ‘eve’ to ‘capu’ edge might be a false negative prediction since ‘eve’ is also known to repress segment polarity genes whereas ‘capu’ is known to have necessary functions in polarity establishment; see Sections ‘Gene Snapshot’ of ‘eve’ and ‘capu’ (http://flybase.org/reports/FBgn0000256). *TGS-Lite+.mf15* misses both ‘mmy’ and ‘capu’.

#### Gene ‘ey’

‘ey’ is known to be a master regulator for eye development [32]. For this gene, an interesting observation is made in the larval stage where *TGS.mf14*, *TGS-Lite.mf15* and *TGS-Lite+.mf15* all agree that ‘ey’ regulates itself. There are hitherto no experimental evidences of protein ‘ey’ binding to the TF-binding site of gene ‘ey’. However, there is an experimental evidence of protein ‘ey’ autoregulating its binding to its target proteins. This autoregulation takes place through interactions of two distinct DNA-binding domains of protein ‘ey’: Paired Domain (PD) and Homeodomain (HD). ‘ey’ PD is essential for the expressions of key TFs in retinal development. Tanaka-Matakatsu et al. [33] suggest that ‘ey’ HD can physically interact with ‘ey’ PD, inhibiting ‘ey’ PD’s ability to bind to its targets. Given that ‘ey’ is a master regulator and autoregulates itself at the Protein-Protein Interaction (PPI) level, the authors hypothesize that ‘ey’ autoregulates itself also at the Protein-DNA Interaction (PDI) level. The experimental verification of this hypothesis presents another novel opportunity for future.

#### Gene ‘prd’

‘prd’ plays a regulatory role in the anterior-posterior segmentation of embryos and gene ‘eve’ is known to be one of its regulatees. Like *TGS.mf14*, *TGS-Lite.mf15* correctly identifies ‘eve’ as a regulatee of ‘prd’ in the embryonic stage. However, *TGS-Lite+.mf15* misses this regulatee. Moreover, *TGS-Lite.mf15* predicts ‘capu’ to be a regulatee of ‘prd’ in the same stage whereas *TGS.mf14* does not make that prediction. Since ‘prd’ and ‘capu’ are known to be essential for developing male and female fertility, respectively, they are likely to have an inhibitory relationship between them; see Section ‘Gene Snapshot’ of ‘prd’ (http://flybase.org/reports/FBgn0003145) and ‘capu’ (http://flybase.org/reports/FBgn0000256). If that is the case, then it is a true positive prediction on *TGS-Lite.mf15*’s part. *TGS-Lite+.mf15* misses this potential regulatee as well. On the other hand, *TGS-Lite.mf15* predicts seven more regulatees than *TGS-Lite+.mf15* in the embryonic stage. No supporting information is found for these regulatees, suggesting that *TGS-Lite+.mf15* correctly rejects these potentially false positive edges.

#### Summary of Findings

This study demonstrates that *TGS-Lite.mf15* predicts potentially more true edges compared to that of *TGS.mf14*. On the other hand, the set of predicted edges by *TGS-Lite+.mf15* is almost a proper subset of that of *TGS-Lite.mf15*, resulting in a more precise prediction with less true positives at the benefit of a less number of potentially false positive edges.

#### Comparison with Alternative Algorithms

{*TVDBN-0, TVDBN-bino-hard, TVDBN-bino-soft*} fail to process any DmLc3 sub-dataset with the given memory. *ARTIVA* succeeds with DmLc3E and fails for other sub-datasets, potentially due to implementation issues. A comparison between {*ARTIVA, TGS-Lite.mf15, TGS-Lite+.mf15*} on DmLc3E reestablishes *TGS-Lite+’s* superiority in terms of speed, and balancing false with true positives (see Section 4.12 of the supplementary document for details).

## 6 Summary and Future Work

In this paper, two novels algorithms, namely *TGS-Lite* and *TGS-Lite+*, are proposed to reconstruct time-varying GRNs underlying a given time series gene expression dataset. It is assumed that the given dataset is complete (no missing values) and has multiple time series. The novelty of the proposed algorithms is that they combine three desired properties – flexibility, time-efficiency and memory-efficiency – in a single framework. Among the prior state-of-the-art algorithms, it is observed that the algorithms that offer state-of-the-art reconstruction power follow flexible frameworks where the ‘smoothly time-varying assumption’ is not enforced on the GRN structures. By employing such a flexible framework, the proposed algorithms are able to offer state-of-the-art reconstruction power in regard to three benchmark datasets. Moreover, they offer such power at state-of-the-art time complexities. It can be noted that there are two prior state-of-the-art algorithms, namely *TGS* and *TGS+*, that provide the same reconstruction power and time complexities. However, their memory-requirements grow exponentially with the number of genes; that of the proposed algorithms grow only linearly.

While both of the proposed algorithms demonstrate state-of-the-art reconstruction power, their strengths lie in different areas. *TGS-Lite* specialises in true positive detection power, making it suitable for applications targeted to discoveries of novel biomarkers. On the other hand, *TGS-Lite+* sacrifices a reasonable amount of true positive detection power to gain a considerable amount of false positive rejection power. Thus, it specializes in overall correctness (F1-score), making it state-of-the-art algorithm for reconstructing time-varying GRNs as correctly as possible and as efficiently as possible. The real-life applicability of *TGS-Lite* and *TGS-Lite+* is demonstrated with a D. melanogaster life cycle dataset. Both the algorithms reconstruct meaningful sequences of time-varying GRNs that help in explaining the developmental process of D. melanogaster through different stages of life.

The flexible framework underlying the proposed algorithms decomposes the reconstruction problem into atomic problems of identifying the regulators of every gene at every time interval. These atomic problems are solved independently of each other without imposing any global constraint, thus providing flexibility. For solving each atomic problem, a two-step learning strategy is deployed. In the first step, a set of candidate regulators is shortlisted for the concerned gene. In the final step, the highest scoring (w.r.t. the BIC scoring function) subset of the shortlisted candidate set is chosen as the final set of regulators for that gene during the corresponding time interval. During this step, every subset is generated in real-time, immediately before its score needs to be calculated, and removed immediately after the score is calculated. Thus, the proposed algorithms achieve a significantly higher memory-efficiency than the prior state-of-the-art algorithms that simultaneously hold all the subsets and their scores in memory. On the other hand, the shortlisting strategy in the first step significantly reduces the time complexity of the second step, thus providing the desired time-efficiency.

Nevertheless, there are opportunities for further improvements. The shortlist of candidate regulators which is prepared for each gene in the first step, is time-invariant. Therefore, the true regulators, which are active across a small number of time intervals, may not get shortlisted. To mitigate this issue, a novel shortlisting strategy can be designed to accommodate time-interval-specific shortlists for each gene. Moreover, the proposed shortlisting strategy requires the data to be discretised, which may incur a loss of information. The same issue is true for the second step which requires discretised data for calculating the BIC scores. Therefore, devising novel shortlisting and scoring strategies that can deal with continuous data remains another challenge. Next challenge lies in enhancing reconstruction power further by integrating time-series gene expression data with auxiliary data, such as knock-out gene expressions [27] and time-series protein expressions [34]. Moreover, recently proposed deep learning-based GRN analysis methods can be explored in the downstream analysis of reconstructed GRNs such as discovering novel biomarkers, functional communities of genes etc. ([35], [36], [37]). Thus, the dedicated pursuit of a more powerful framework is driven by the hope that its marriage with high-throughput measurement techniques will pave the way for a better understanding of temporal biological processes such as development and pathogenesis.

## ACKNOWLEDGEMENTS

The authors acknowledge the Department of Biotechnology, Govt of India for the financial support for the project BT/COE/34/SP28408/2018, and IITG for MHRD Fellowships to SP.

---

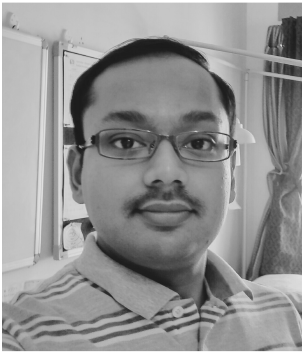

**Saptarshi Pyne** is a PhD student in Dr Ashish Anand’s research group. His research area is temporal progression modelling of biological systems. Saptarshi believes in a future where biomolecular signals are measured in-vivo and analysed in near real time.

**Website:** http://sap01.github.io/

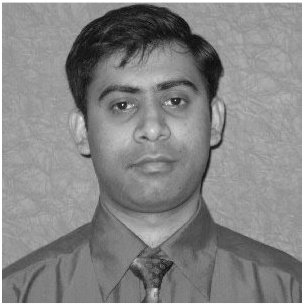

**Ashish Anand** is an associate professor at the Department of Computer Science and Engineering, Indian Institute of Technology Guwahati (IITG), India. Earlier, he was a member of the European Consortium, BaSySBio at the Systems Biology Lab, Institut Pasteur, Paris. His main research area is temporal progression modelling of biological systems.

**Website:** http://iitg.ac.in/anand.ashish/

## Footnotes

1. ‘Fan-in’ of a node represents the number of regulators the node has. The max fan-in restriction puts an upper bound on how many regulators a node can have.

