## Supplementary document for "Rapid Reconstruction of Time-varying Gene Regulatory Networks with Limited Main Memory"

### Supplementary Document for the Paper Titled ‘Rapid Reconstruction of Time-varying Gene Regulatory Networks with Limited Main Memory’

By

**Saptarshi Pyne and Ashish Anand**

**August 29, 2019**

### Chapter 1

### Datasets

The files mentioned in this chapter can be found at: `https://github.com/aaiitg-grp/TGS-tcbb/tree/master/datasets` . Please note that, when notation '$(x, y)$' is used in the context of a cell in a matrix, x and y represent the row and the column of the cell, respectively.

#### 1.1 Synthetic DREAM3 In Silico Network Inference Challenge Datasets

### 1.1.1 Ds10n

File: 'InSilicoSize10-Yeast1-trajectories.tsv'. The columns represent the genes and the rows represent the time points. There are a total of 10 genes identified by $\{G1, G2, \ldots, G10\}$ and 21 distinct time points denoted by $\{0.0, 10.0, 20.0, \ldots, 200.0\}$. Ds10n contains 4 separate time series. Therefore, there will be 4 separate rows for each pair of (time point ID, gene ID), which represent the expression of the gene at the same time point in different time series. The wild type (WT) values for the genes can be found at the 'wt' row of file 'InSilicoSize10-Yeast1-null-mutants.tsv'.

The true network file: 'DREAM3GoldStandard_InSilicoSize10_Yeast1_TrueNet.RData'. Please start a R session in the same directory where you have saved the file and load the RData file as shown below.

```
> load('DREAM3GoldStandard_InSilicoSize10_Yeast1_TrueNet.RData')
> ls() ## list objects in the current workspace
[1] "true.net.adj.matrix"
```

The 'true.net.adj.matrix' R object is the adjacency matrix of the true network. It is a binary matrix of dimension $(10 \times 10)$. Here, $(G3, G1) = 1$ implies that there is a directed edge from $G3$ to $G1$ in the true network. On the other hand, $(G3, G2) = 0$ implies that there does not exist any directed edge from $G3$ to $G2$ in the true network.

### 1.1.2 Ds50n

File: 'InSilicoSize50-Yeast1-trajectories.tsv'. The columns represent the genes and the rows represent the time points. There are a total of 50 genes identified by $\{G1, G2, \ldots, G50\}$ and 21 distinct time points denoted by $\{0.0, 10.0, 20.0, \ldots, 200.0\}$. Ds50n contains 23 separate time series. Therefore, there will be 23 separate rows for each pair of (time point ID, gene ID), which represent the expression of the gene at the same time point in different time series. The wild type (WT) values for the genes can be found at the 'wt' row of file 'InSilicoSize50-Yeast1-null-mutants.tsv'.

The true network file: 'DREAM3GoldStandard_InSilicoSize50_Yeast1_TrueNet.RData'. Please start a R session in the same directory where you have saved the file and load the RData file as shown below.

```
> load('DREAM3GoldStandard_InSilicoSize50_Yeast1_TrueNet.RData')
> ls() ## list objects in the current workspace
[1] "true.net.adj.matrix"
```

The 'true.net.adj.matrix' R object is the adjacency matrix of the true network. It is a binary matrix of dimension $(50 \times 50)$. Here, $(G2, G1) = 1$ implies that there is a directed edge from $G2$ to $G1$ in the true network. On the other hand, $(G2, G4) = 0$ implies that there does not exist any directed edge from $G2$ to $G4$ in the true network.

### 1.1.3 Ds100n

File: 'InSilicoSize100-Yeast1-trajectories.tsv'. The columns represent the genes and the rows represent the time points. There are a total of 100 genes identified by $\{G1, G2, \ldots, G100\}$ and 21 distinct time points denoted by $\{0.0, 10.0, 20.0, \ldots, 200.0\}$. Ds100n contains 46 separate time series. Therefore, there will be 46 separate rows for each pair of (time point ID, gene ID), which represent the expression of the gene at the same time point in different time series. The wild type (WT) values for the genes can be found at the 'wt' row of file 'InSilicoSize100-Yeast1-null-mutants.tsv'.

The true network file: 'DREAM3GoldStandard_InSilicoSize100_Yeast1_TrueNet.RData'. Please start a R session in the same directory where you have saved the file and load the RData file as shown below.

```
> load('DREAM3GoldStandard_InSilicoSize100_Yeast1_TrueNet.RData')
> ls() ## list objects in the current workspace
[1] "true.net.adj.matrix"
```

The 'true.net.adj.matrix' R object is the adjacency matrix of the true network. It is a binary matrix of dimension $(100 \times 100)$. Here, $(G2, G3) = 1$ implies that there is a directed edge from $G2$ to $G3$ in the true network. On the other hand, $(G2, G1) = 0$ implies that there does not exist any directed edge from $G2$ to $G1$ in the true network.

#### 1.2 Real Drosophila melanogaster (Dm) Life Cycle Dataset (DmLc)

##### 1.2.1 DmLc3E

File: 'DmLc3E.RData'. Please start a R session in the same directory where you have saved the file and load the RData file as shown below.

```
> load('DmLc3E.RData')
> ls()
[1] "input.data"
```

The 'input.data' R object is a data matrix of dimension $(30 \times 588)$. The rows and columns represent the time points and the genes, respectively. The column names are the gene names.

##### 1.2.2 DmLc3L

File: 'DmLc3L.RData'. Please start a R session in the same directory where you have saved the file and load the RData file as shown below.

```
> load('DmLc3L.RData')
> ls()
[1] "input.data"
```

The 'input.data' R object is a data matrix of dimension $(10 \times 588)$. The rows and columns represent the time points and the genes, respectively. The column names are the gene names.

##### 1.2.3 DmLc3P

File: 'DmLc3P.RData'. Please start a R session in the same directory where you have saved the file and load the RData file as shown below.

```
> load('DmLc3P.RData')
> ls()
[1] "input.data"
```

The ‘input.data’ R object is a data matrix of dimension ($18 \times 588$). The rows and columns represent the time points and the genes, respectively. The column names are the gene names.

##### 1.2.4 DmLc3A

File: ‘DmLc3A.RData’. Please start a R session in the same directory where you have saved the file and load the RData file as shown below.

```
> load('DmLc3A.RData')
> ls()
[1] "input.data"
```

The ‘input.data’ R object is a data matrix of dimension ($8 \times 588$). The rows and columns represent the time points and the genes, respectively. The column names are the gene names.

### Chapter 2

### Source Code

The first step towards reproducing the results is replication of the experimental environment. It can be achieved by completing the following sub-steps:

- Install the same hardware
- Install the same Operating System (OS) with the same version
- Install the same version(s) of the required programming language(s)
- Execute the same implementations

Please find the hardware configuration and OS details of the experimental environment in Section 2.2. It might be infeasible to purchase the same hardware. In that case, an available hardware can be used. Although, it needs to be noted that a change in the hardware might cause the runtime to vary; a smaller main memory may even cause a memory inadequacy related error.

On the other hand, installing the same OS with the same version is easier than replicating the hardware. However, if you wish to carry on with a different OS or a different version of the same OS, some changes might be necessary as will discussed in this chapter. Since, the implementations are in the R programming language, the amount of necessary changes is expected not to be overwhelming.

The next step is to install the same version(s) of the required programming language(s). Both *TGS-Lite* and *TGS-Lite+* are implemented in the R programming language [8] version 3.5.1. How this specific R version is installed in the experimental environment is described in Section 2.3.

Finally, how to re-use the implementations of the *TGS-Lite* and *TGS-Lite+* algorithms is demonstrated in Section 2.4.

#### 2.1 Notations

- When notation '$(x, y)$' is used in the context of a cell in a matrix, x and y represent the row and the column of the cell, respectively.
- 'pack1::func1()' represents the 'func1()' function of R package 'pack1'.

#### 2.2 Hardware Configuration of the Experimental Environment

Experiments are performed on an Intel® computing server with the following configuration:

- Architecture: x86_64
- CPUs: Two Intel® Xeon® X5675 @ 3.07GHz CPUs
- Main Memory: 31 GB

- Swap Space: 34 GB
- Cache: {L1d cache: 32 KB, L1i cache: 32 KB, L2 cache: 256 KB, L3 cache: 12288 KB}
- Secondary Storage: 4.1 TB
- Operating System: Ubuntu 12.04.5 LTS (Codename: Precise)

#### 2.3 Installing R version 3.5.1 in the Experimental Environment

Please refer to: `https://github.com/sap01/TGS-Lite-supplem/blob/master/README.md` .

#### 2.4 Executing the *TGS-Lite* and *TGS-Lite+* Algorithms in the Experimental Environment

The R implementations of the *TGS-Lite* and *TGS-Lite+* algorithms are saved as a *R* project tarball ( `https://github.com/sap01/TGS-Lite-supplem/blob/master/sourcecode/TGS-Lite-2019-04-15.tar.gz` ). Copy this file to '/home/saptarshi/R/R-3.5.1/projects':

```
$ mv TGS-Lite-2019-04-15.tar.gz /home/saptarshi/R/R-3.5.1/projects
```

After that, unbundle the project using 'packrat':

```
%% Go to the project directory
$ cd /home/saptarshi/R/R-3.5.1/projects

%% Open a R prompt
$ R351

## Set your favourite CRAN repo, e.g.,
## https://cran.rstudio.com/
> options(repos=structure(c(CRAN="https://cran.rstudio.com/")))

## Attach 'packrat' package
> library(packrat)

## Unbundle the project inside the current directory
> packrat::unbundle('TGS-Lite-2019-04-15.tar.gz', getwd())
```

Once unbundled, a new project directory will be created with name 'TGS-Lite'. Go inside the project directory:

```
%% Go to /home/saptarshi/R/R-3.5.1/projects/TGS-Lite
$ cd TGS-Lite
```

Directory 'TGS-Lite' contains all required R scripts and two sub-directories: 'packrat' and 'asset'. The 'packrat' sub-directory is for internal management of 'packrat' and not to be interfered with. The 'asset' sub-directory is the place where the input and the output files are stored. Copy all the dataset files inside this sub-directory. For example, let us assume that the directory corresponding to `https://github.com/aaiitg-grp/TGS/tree/master/datasets` , in your local computer, is '/home/saptarshi/datasets'. Then copy all the files from that directory to 'TGS-Lite/asset':

```
$ scp /home/saptarshi/datasets/* asset
```

The driver R script is 'TGS-Lite/TGS.R'. It takes the user-defined input parameters in a JSON file format [2]. The JSON file must reside in the 'asset' sub-directory. 'TGS.R' can be executed with the following command:

```
%% Assuming 'TGS-Lite/asset/input.json' contains the user-defined parameters.
%% Note that only 'input.json' is used instead of 'asset/input.json'. This
%% is because 'TGS.R' is programmed to search for the input JSON files in
%% the 'asset' sub-directory.
%% The '&' symbol instructs the execution process to start in the background.
$ nohup time /home/saptarshi/R/R-3.5.1/bin/Rscript TGS.R input.json &

%% Prints the process ID
[1] 8172
```

Since, the execution process is performed in the background, the bash command prompt can be used for other tasks. Once the execution is complete, a 'Done' message is automatically displayed in bash. However, if you wish to monitor the execution of the process, you may do so with the 'top' command as shown below:

```
%% Show details of the process with ID 8172
$ top -p 8172
```

The input JSON files required for reproducing each and every result are already stored in the 'asset' sub-directory. Please choose the appropriate input JSON file from Table 2.1 for your desired experiment. Each JSON file is a collection of one or more {name, value} pairs. Each such pair corresponds to the name and the value of a specific parameter. The parameters in an input JSON file are described in Section 2.4.1.

Table 2.1: Description of input JSON files in the '/TGS-Lite/asset' directory

| **JSON Filename** | **Algorithm** | **Target Experiment** |
|---|---|---|
| input.Ds10n.2L.wt.aro1.mi.pca.cmi.CLR.lite.bic.mf14.json | TGS-Lite | Dataset Ds10n with *2L.wt* discretization |
| input.Ds10n.2L.wt.aro1.mi.pca.cmi.CLR.lite.bic.mf14.p10.json | TGS-Lite.p10 | Dataset Ds10n with *2L.wt* discretization |
| input.Ds50n.2L.wt.aro1.mi.pca.cmi.CLR.lite.bic.mf14.json | TGS-Lite | Dataset Ds50n with *2L.wt* discretization |
| input.Ds50n.2L.wt.aro1.mi.pca.cmi.CLR.lite.bic.mf14.p10.json | TGS-Lite.p10 | Dataset Ds50n with *2L.wt* discretization |

Continued on next page

**Table 2.1 – continued from previous page**

| JSON Filename | Algorithm | Target Experiment |
|---|---|---|
| input.Ds100n.2L.wt.aro1.mi.pca.cmi.CLR.lite.bic.mf14.json | TGS-Lite | Dataset Ds100n with *2L.wt* discretization, number of cores = 1, max fan-in = 14 |
| input.Ds100n.2L.wt.aro1.mi.pca.cmi.CLR.lite.bic.mf14.p10.json | TGS-Lite.p10 | Dataset Ds100n with *2L.wt* discretization, number of cores = 10, max fan-in = 14 |
| input.Ds100n.2L.wt.aro1.mi.pca.cmi.CLR.lite.bic.mf14.p3.json | TGS-Lite.mf14 | Dataset Ds100n with *2L.wt* discretization, number of cores = 3, max fan-in = 14 |
| input.Ds100n.2L.wt.aro1.mi.pca.cmi.CLR.lite.bic.mf14.p7.json | TGS-Lite.mf14 | Dataset Ds100n with *2L.wt* discretization, number of cores = 7, max fan-in = 14 |
| input.Ds100n.2L.wt.aro1.mi.pca.cmi.CLR.lite.bic.mf15.p10.json | TGS-Lite.p10 | Dataset Ds100n with *2L.wt* discretization, number of cores = 10, max fan-in = 15 |

Continued on next page

**Table 2.1 – continued from previous page**

| JSON Filename | Algorithm | Target Experiment |
|---|---|---|
| input.Ds100n.2L.wt.aro1.mi.pca.cmi.CLR.lite.bic.mf16.p10.json | TGS-Lite.p10 | Dataset Ds100n with *2L.wt* discretization, number of cores = 10, max fan-in = 16 |
| input.Ds100n.2L.wt.aro1.mi.pca.cmi.CLR.lite.bic.mf17.p10.json | TGS-Lite.p10 | Dataset Ds100n with *2L.wt* discretization, number of cores = 10, max fan-in = 17 |
| input.Ds100n.2L.wt.aro1.mi.pca.cmi.CLR.lite.bic.mf18.p10.json | TGS-Lite.p10 | Dataset Ds100n with *2L.wt* discretization, number of cores = 10, max fan-in = 18 |
| input.Ds10n.2L.wt.aro1.mi.pca.cmi.aracne.CLR.lite.bic.mf14.json | TGS-Lite+ | Dataset Ds10n with *2L.wt* discretization |
| input.Ds10n.2L.wt.aro1.mi.pca.cmi.aracne.CLR.lite.bic.mf14.p10.json | TGS-Lite+.p10 | Dataset Ds10n with *2L.wt* discretization |
| input.Ds50n.2L.wt.aro1.mi.pca.cmi.aracne.CLR.lite.bic.mf14.json | TGS-Lite+ | Dataset Ds50n with *2L.wt* discretization |
| input.Ds50n.2L.wt.aro1.mi.pca.cmi.aracne.CLR.lite.bic.mf14.p10.json | TGS-Lite+.p10 | Dataset Ds50n with *2L.wt* discretization |

Continued on next page

**Table 2.1 – continued from previous page**

| JSON Filename | Algorithm | Target Experiment |
|---|---|---|
| input.Ds100n.2L.wt.aro1.mi.pca.cmi.aracne.CLR.lite.bic.mf14.json | TGS-Lite+ | Dataset Ds100n with *2L.wt* discretization, number of cores = 1, max fan-in = 14 |
| input.Ds100n.2L.wt.aro1.mi.pca.cmi.aracne.CLR.lite.bic.mf14.p10.json | TGS-Lite+.p10 | Dataset Ds100n with *2L.wt* discretization, number of cores = 10, max fan-in = 14 |
| input.Ds100n.2L.wt.aro1.mi.pca.cmi.aracne.CLR.lite.bic.mf14.p3.json | TGS-Lite+.mf14 | Dataset Ds100n with *2L.wt* discretization, number of cores = 3, max fan-in = 14 |
| input.Ds100n.2L.wt.aro1.mi.pca.cmi.aracne.CLR.lite.bic.mf14.p7.json | TGS-Lite+.mf14 | Dataset Ds100n with *2L.wt* discretization, number of cores = 7, max fan-in = 14 |
| input.Ds100n.2L.wt.aro1.mi.pca.cmi.aracne.CLR.lite.bic.mf15.p10.json | TGS-Lite+.p10 | Dataset Ds100n with *2L.wt* discretization, number of cores = 10, max fan-in = 15 |

Continued on next page

**Table 2.1 – continued from previous page**

| **JSON Filename** | **Algorithm** | **Target Experiment** |
|---|---|---|
| input.Ds100n.2L.wt.aro1.mi.pca.cmi.aracne.CLR.lite.bic.mf16.p10.json | TGS-Lite+.p10 | Dataset Ds100n with *2L.wt* discretization, number of cores = 10, max fan-in = 16 |
| input.Ds100n.2L.wt.aro1.mi.pca.cmi.aracne.CLR.lite.bic.mf17.p10.json | TGS-Lite+.p10 | Dataset Ds100n with *2L.wt* discretization, number of cores = 10, max fan-in = 17 |
| input.Ds100n.2L.wt.aro1.mi.pca.cmi.aracne.CLR.lite.bic.mf18.p10.json | TGS-Lite+.p10 | Dataset Ds100n with *2L.wt* discretization, number of cores = 10, max fan-in = 18 |
| input.DmLc3E.aro1.mi.pca.cmi.CLR.lite.bic.mf15.p10.json | TGS-Lite.mf15 | Dataset DmLc3E, number of cores = 10, max fan-in = 15 |
| input.DmLc3L.aro1.mi.pca.cmi.CLR.lite.bic.mf15.p10.json | TGS-Lite.mf15 | Dataset DmLc3L, number of cores = 10, max fan-in = 15 |
| input.DmLc3P.aro1.mi.pca.cmi.CLR.lite.bic.mf15.p10.json | TGS-Lite.mf15 | Dataset DmLc3P, number of cores = 10, max fan-in = 15 |

Continued on next page

**Table 2.1 – continued from previous page**

| JSON Filename | Algorithm | Target Experiment |
|---|---|---|
| input.DmLc3A.aro1.mi.pca.cmi.CLR.lite.bic.mf15.p10.json | TGS-Lite.mf15 | Dataset DmLc3A, number of cores = 10, max fan-in = 15 |
| input.DmLc3E.aro1.mi.pca.cmi.aracne.CLR.lite.bic.mf15.p10.json | TGS-Lite+.mf15 | Dataset DmLc3E, number of cores = 10, max fan-in = 15 |
| input.DmLc3L.aro1.mi.pca.cmi.aracne.CLR.lite.bic.mf15.p10.json | TGS-Lite+.mf15 | Dataset DmLc3L, number of cores = 10, max fan-in = 15 |
| input.DmLc3P.aro1.mi.pca.cmi.aracne.CLR.lite.bic.mf15.p10.json | TGS-Lite+.mf15 | Dataset DmLc3P, number of cores = 10, max fan-in = 15 |
| input.DmLc3A.aro1.mi.pca.cmi.aracne.CLR.lite.bic.mf15.p10.json | TGS-Lite+.mf15 | Dataset DmLc3A, number of cores = 10, max fan-in = 15 |

Once 'TGS.R' completes its execution, a file named 'nohup.out' would be created inside the 'TGS-Lite' directory and all the output files would be saved in an output directory inside 'TGS-Lite/asset'. In order to find the name of the output directory, please open 'nohup.out' in a text editor. There would be a line stating "The output directory name is:"; its next line would reveal the output directory name. For example:

```
[1] "The output directory name is:"
[1] "/home/saptarshi/R/R-3.5.1/projects/TGS-Lite/asset/output20190103161755"
```

The output directory name is 'output' followed by the current timestamp in the 'YYYYmmddHHMMSS' format, as given by the R command 'format(Sys.time(), "%Y%m%d%H%M%S")'. Once the name is known, please go to the directory:

```
$ cd asset/output20190103161755

%% Also move the 'nohup.out' to the output directory.
%% Otherwise, the next execution of 'TGS.R' would
```

```
%% overwrite it.
$ mv ../../nohup.out .
```

Now, the output directory contains all the output files for this particular experiment. Section 2.4.2 describes how to interpret these files.

##### 2.4.1 Descriptions of Parameters in an Input JSON File

**input.data.filename** The 'input.data.filename' parameter can have filenames with either the '.tsv' or the '.RData' extension. If the file has the '.tsv' extension, then the first column should contain the time point IDs except the $(1, 1)^{th}$ cell. The first row should contain the gene names, except the $(1, 1)^{th}$ cell. The $(1, 1)^{th}$ cell does not carry any meaning. If the file has the '.RData' extension, then the underlying object must have the name 'input.data'. In 'input.data', the column names and the row names represent the gene names and the time point IDs, respectively. For either of '.tsv' or '.RData' input, Multiple rows with the same time point ID represent multiple replicates at the same time point. In other words, those rows belong to the same time point but at different time series. The time points belonging to the same time series must be together and in ascending order. An exemplary dataset with three genes {G1, G2, G3}, two time points {t1, t2} and two time series is shown below.

```
Time  G1           G2           G3
t1    0.8272480342 0.7257430901 0.3894130418
t2    0.6542518342 0.6470658823 0.5088904888
t1    0.3519554463 0.3551279726 0.3207993604
t2    0.4871730974 0.3706990326 0.447523615
```

**num.timepts** The 'num.timepts' parameter defines the number of distinct time points in the input data file.

**true.net.filename** If 'true.net.filename' is an empty string, then it is implied that the true rolled network is not known a prior. But if it is not an empty string, then it must be a '.RData' file. In that case, the underlying object must be of name 'true.net.adj.matrix'. In 'true.net.adj.matrix', the row names and the column names represent the gene names. It is a binary matrix. If $(i, j)^{th}$ cell contains 1, then there exists a directed edge from the $i^{th}$ gene to the $j^{th}$ gene in the true network. Else if $(i, j)^{th}$ cell contains 0, then that edge does not exist in the true network.

**input.wt.data.filename** The 'input.wt.data.filename' parameter provides the name of the file which has the Wild Type (WT) values of the genes. If its value is an empty string, then the WT values of the genes are not known. Otherwise, it must be a file with '.tsv' extension. Inside the file, the first row should contain the gene names, except the $(1, 1)^{th}$ cell. Then the second row should contain the WT values of those genes, except the $(2, 1)^{th}$ cell. Rest of the file does not carry any meaning.

**is.discrete** The 'is.discrete' parameter takes value 'true' or 'false', depending on whether the input data is already discretized or not, respectively.

**num.discr.levels** The 'num.discr.levels' parameter provides the number of discrete levels each gene has if the input dataset is already discretized ("is.discrete": true). On the other hand, 'num.discr.levels' provides the number of discrete levels into which each gene needs to be discretized if the input dataset is not yet discretized ("is.discrete": false).

**discr.algo** The 'discr.algo' parameter is used to specify the name of the discretization algorithm to be used in case the input data needs to be discretized. For the time being, only two options are available:

‘discretizeData.2L.wt.l’ and ‘discretizeData.2L.Tesla’, which represent algorithms *2L.wt* and *2L.Tesla*, respectively [7].

**mi.estimator** The ‘mi.estimator’ parameter specifies which method to use for estimating the mutual information matrix. For all the experiments, the ‘mi.pca.cmi’ estimator is used. The original source code of ‘mi.pca.cmi’ is written in MATLAB and available as function ‘cmi()’ at `http://www.comp-sysbio.org/grn/pca_cmi.m`.

**apply.aracne** ‘apply.aracne’ is a boolean parameter. The *ARACNE* step is applied to refine the raw mutual information matrix only if this parameter is set to ‘true’. If ‘apply.aracne’ is set to ‘false’, then the *ARACNE* step is excluded and the raw mutual information matrix is used for further computation. Please see the description of parameter ‘use.lite’ below to understand how it is combined with parameter ‘apply.aracne’ to choose different algorithms.

**clr.algo** This parameter indicates which *CLR* variant to employ. Currently, only one variant is available, namely ‘CLR’.

**max.fanin** The ’max.fanin’ parameter provides the maximum number of regulators each gene can have.

**allow.self.loop** The ‘allow.self.loop’ parameter takes value ‘true’ or ‘false’, depending on whether to allow self loops in the predicted rolled network or not.

**use.lite** The ‘use.lite’ parameter takes value ‘true’ or ‘false’. Depending upon the combination of values set to parameters ‘apply.aracne’ and ‘use.lite’, you will be able to choose different algorithms (Table 2.2 ).

Table 2.2: How to choose different algorithms by setting different values to parameters ‘apply.aracne’ and ‘use.lite’

| **apply.aracne** | **use.lite** | **Algorithm** |
|---|---|---|
| false | false | *TGS* |
| false | true | *TGS-Lite* |
| true | false | *TGS+* |
| true | true | *TGS-Lite+* |

**scoring.func** The ‘scoring.func’ parameter indicates the scoring function to be used for scoring different Bayesian network structures and thus learning the highest scoring structure. For the time being, this parameter can only take the value ‘BIC’.

**parallel** Please set the ‘parallel’ parameter to ‘true’ if you wish to utilize multiple computing cores in parallel; otherwise, set it to ‘false’ so that the execution is performed in a serial fashion using a single core.

**max.num.cores** When the ‘parallel’ parameter is set to ‘true’, the ‘max.num.cores’ parameter can be used to set an upper limit to how many computing cores the R script should use. In that case, the script actually uses the number of cores which equals to min(the value of ‘max.num.cores’, number of genes, (available number of computing cores - 1)). The subtraction by one in the last expression is to ensure that at least one core is left for monitoring purposes. On the other hand, when the ‘parallel’ parameter is set to ‘false’, the ‘max.num.cores’ parameter takes its default value ‘1’.

##### 2.4.2 Description of *TGS* Output Files

**output.txt** This file saves the console output generated by 'TGS.R' and its callee R scripts. Please open it in a text editor. At the very beginning, there is a line stating 'elapsed.time just after CLR step=' followed by the time taken by the *CLR* step in seconds. For example:

```
elapsed.time just after CLR step=

user  system elapsed
0.000   0.000   0.003
```

In this case, the 'elapsed' time is considered as the completion time for the *CLR* step, which is 0.003 seconds.

At the very end of the file, there is a line stating 'elapsed.time =' followed by the time taken by the chosen algorithm in seconds. For example:

```
elapsed.time =

user  system elapsed
5.684   0.100   5.789
```

In this case, the 'elapsed' time is considered as the runtime of the chosen algorithm, which is 5.789 seconds.

Only for the experiments where the true rolled network is provided as an input for evaluating the learning power of *TGS*, there are four lines just before the 'elapsed.time =' line. Among these four lines, the first line states 'Result TGS vs True =' and the second line is a blank line; finally, the last two lines depict the evaluation metrics of *TGS*. For example:

```
Result TGS vs True =

     TP TN FP FN TPR       FPR       FDR       PPV  ACC       MCC         F
[1,]  3 80 10  7 0.3 0.1111111 0.7692308 0.2307692 0.83 0.1684986 0.2608696
```

In this case, the metrics are read as 'TP = 3', 'TN = 80' and so on.

The rest of the lines in the file are for debugging purposes only.

**di.net.adj.matrix.RData and net.sif** File 'di.net.adj.matrix.RData' contains the predicted rolled network's adjacency matrix. The matrix can be analysed by loading the file in a R session:

```
## Load object 'di.net.adj.matrix'
> load('di.net.adj.matrix.RData')
```

The row names and the column names correspond to the input gene names. The $(i, j)^{th}$ cell takes value 1 if and only if there is a directed edge from the $i^{th}$ gene to the $j^{th}$ gene in the predicted network; otherwise, it takes value 0.

'net.sif' is the Cytoscape compatible SIF format of 'di.net.adj.matrix.RData'.

**unrolled.DBN.adj.matrix.list.RData** This file contains the unrolled time-varying Gene Regulatory Networks. The networks can be analysed by loading the file in a R session:

```
## Load object 'unrolled.DBN.adj.matrix.list'
> load('unrolled.DBN.adj.matrix.list.RData')
```

The object 'unrolled.DBN.adj.matrix.list' is a list of $(T - 1)$ matrices, where $T$ denotes the total number of time points in the input dataset. The $t^{th}$ element of the list is the adjacency matrix of the network that represents the predicted gene regulations during the time interval between time points $t$ and $(t + 1)$. For example, when $T = 21$:

```
## Length of the list = (T - 1)
> length(unrolled.DBN.adj.matrix.list)
[1] 20

## Print the first network i.e.
## predicted gene regulations during
## the time interval between the 1st
## and the 2nd time points
> unrolled.DBN.adj.matrix.list[[1]]
    v1 v2 v3 v4 v5 v6 v7 v8 v9 v10
v1   0  0  0  0  0  0  0  0  0   0
v2   1  0  0  1  0  0  0  0  0   0
v3   0  0  0  0  0  0  0  0  0   0
v4   0  1  0  0  0  0  0  0  0   0
v5   0  0  0  0  0  0  0  0  0   0
v6   0  0  0  0  0  0  0  0  0   0
v7   0  0  0  0  0  0  0  0  0   0
v8   0  0  0  0  0  0  0  0  0   0
v9   0  0  0  0  0  0  0  0  0   0
v10  0  0  0  0  0  0  0  0  0   0
```

It needs to be noted that each original gene name is replaced with 'v' followed by its index. For example, the original gene names in dataset Ds10n are {G1, G2, ..., G10} in the given order. Therefore, they are replaced with {v1, v2, ..., v10}. Such name replacement strategy is performed for all intermediate outputs (i.e. all outputs but the final output 'di.net.adj.matrix.RData') to avoid unexpected symbols in the original gene names that might cause errors. If required, the original names can be restored with the help of 'di.net.adj.matrix.RData':

```
## Load object 'di.net.adj.matrix'
> load('di.net.adj.matrix.RData')

## List objects in the current workspace
> ls()
[1] "di.net.adj.matrix"             "unrolled.DBN.adj.matrix.list"

## Get the original gene names
> orig.names <- colnames(di.net.adj.matrix)
> orig.names
[1] G1"  "G2"  "G3"  "G4"  "G5"  "G6"  "G7"  "G8"  "G9"  "G10"

## Restore the original gene names
> rownames(unrolled.DBN.adj.matrix.list[[1]]) <- orig.names
> colnames(unrolled.DBN.adj.matrix.list[[1]]) <- orig.names

## Print the first network
> unrolled.DBN.adj.matrix.list[[1]]
    G1 G2 G3 G4 G5 G6 G7 G8 G9 G10
G1   0  0  0  0  0  0  0  0  0   0
G2   1  0  0  1  0  0  0  0  0   0
G3   0  0  0  0  0  0  0  0  0   0
G4   0  1  0  0  0  0  0  0  0   0
G5   0  0  0  0  0  0  0  0  0   0
G6   0  0  0  0  0  0  0  0  0   0
G7   0  0  0  0  0  0  0  0  0   0
G8   0  0  0  0  0  0  0  0  0   0
```

```
G9   0  0  0  0  0  0  0  0  0   0
G10  0  0  0  0  0  0  0  0  0   0
```

In each predicted time-varying network, the $(i,j)^{th}$ cell takes value 1 if and only if the $i^{th}$ gene regulates the $j^{th}$ gene during that time interval. The predicted rolled network is the union of all the predicted time-varying networks; in other words, the $(i,j)^{th}$ cell in the rolled network adjacency matrix takes value 1 if and only if there exists at least one time-varying network whose adjacency matrix's $(i,j)^{th}$ cell contains value 1, otherwise the former takes value 0.

**mut.info.matrix.RData, mut.info.matrix.pre.aracne.RData, mut.info.matrix.post.aracne.RData, mi.net.adj.matrix.wt.RData and mi.net.adj.matrix.RData** These files save intermediate outputs during the *CLR* step. When 'apply.aracne' is set to false, the raw mutual information matrix is saved in 'mut.info.matrix.RData'. When 'apply.aracne' is set to true, the raw mutual information matrix is saved in 'mut.info.matrix.pre.aracne.RData'. In the latter case, the raw mutual information matrix is then refined by passing it through *ARACNE*. The refined mutual information matrix is saved in 'mut.info.matrix.post.aracne.RData'.

*CLR* takes the raw mutual information matrix (when 'apply.aracne' is false) or the refined mutual information matrix (when 'apply.aracne' is true) as input and produces a weighted *CLR* network adjacency matrix 'mi.net.adj.matrix.wt' (stored in 'mi.net.adj.matrix.wt.RData') where the $(i,j)^{th}$ cell contains a non-zero value if and only if an undirected edge exists between the $i^{th}$ and the $j^{th}$ genes; otherwise, it contains value 0. This matrix is used to compute an unweighted undirected network adjecncy matrix 'mi.net.adj.matrix' (stored in 'mi.net.adj.matrix.RData') by retaining only the top 'max.fanin' (see Paragraph 'max.fanin', Section 2.4.1) number of neighbours for each gene based on the edge weights. In this matrix, the $(i,j)^{th}$ cell contains value 1 if and only if an undirected edge exists between the $i^{th}$ and the $j^{th}$ genes; otherwise, it contains value 0.

**input.data.discr.RData** This file is generated only if the input dataset is not discretized already. In that case, this file represents the input dataset after discretization with the user-defined discretization settings. Please load this file in a R session to analyze the undelying R object 'input.data.discr':

```
## Load a R object named 'input.data.discr'
> load('input.data.discr.RData')
```

'input.data.discr' is a data matrix with the $(i,j)^{th}$ cell representing the discretized expression level of the $j^{th}$ gene in the $i^{th}$ observation of the original input dataset.

**sessionInfo.txt** This file provides the R session information during a specific experiment, as generated by the R function 'sessionInfo()'.

### Chapter 3

### Results

The files mentioned in this chapter can be found at: `https://github.com/sap01/TGS-Lite-supplem/tree/master/results` .

#### 3.1 Results of Algorithms *TGS-Lite* and *TGS-Lite+*

The names of the output directories for different experimental conditions (represented by distinct input JSON files) of the *TGS-Lite* and *TGS-Lite+* algorithms are mentioned in Table 3.1. The experimental condition to input JSON file mapping can be found at Table 2.1.

Table 3.1: The names of the output directories for different experimental conditions (represented by distinct input JSON files) of the *TGS-Lite* and *TGS-Lite+* algorithms are mentioned in Table 3.1. The experimental condition to input JSON file mapping can be found at Table 2.1.
For output directories marked with asterisks, the original directory is renamed to ease identification. For example, the original output directory 'output20180828124106' is renamed to 'TGS-Lite-DmLc3E-output20180828124106', which implies that the directory contains the output of the *TGS-Lite* algorithm with the DmLc3E dataset.
The complete set of findings for the TRANSFAC analyses of the predicted GRNs by the *TGS-Lite.mf15* and *TGS-Lite+.mf15* algorithms are presented in 'DmLc3.flybase.ids.redfly.tfs.xlsx'.

| **Input JSON File** | **Output Directory** |
|---|---|
| input.Ds10n.2L.wt.aro1.mi.pca.cmi.CLR.lite.bic.mf14.json | output20180816153824 |
| input.Ds10n.2L.wt.aro1.mi.pca.cmi.CLR.lite.bic.mf14.p10.json | output20190103161755 |
| input.Ds50n.2L.wt.aro1.mi.pca.cmi.CLR.lite.bic.mf14.json | output20190104150028 |
| input.Ds50n.2L.wt.aro1.mi.pca.cmi.CLR.lite.bic.mf14.p10.json | output20181010164834 |
| input.Ds100n.2L.wt.aro1.mi.pca.cmi.CLR.lite.bic.mf14.json | output20190103173707 |
| input.Ds100n.2L.wt.aro1.mi.pca.cmi.CLR.lite.bic.mf14.p10.json | output20181004155044 |
| input.Ds100n.2L.wt.aro1.mi.pca.cmi.CLR.lite.bic.mf14.p3.json | output20190104193753 |
| input.Ds100n.2L.wt.aro1.mi.pca.cmi.CLR.lite.bic.mf14.p7.json | output20190107153805 |
| input.Ds100n.2L.wt.aro1.mi.pca.cmi.CLR.lite.bic.mf15.p10.json | output20180913174859 |
| input.Ds100n.2L.wt.aro1.mi.pca.cmi.CLR.lite.bic.mf16.p10.json | output20180913194702 |
| input.Ds100n.2L.wt.aro1.mi.pca.cmi.CLR.lite.bic.mf17.p10.json | output20180913235624 |
| input.Ds100n.2L.wt.aro1.mi.pca.cmi.CLR.lite.bic.mf18.p10.json | output20180914084121 |
| input.Ds10n.2L.wt.aro1.mi.pca.cmi.aracne.CLR.lite.bic.mf14.json | output20180816152739 |
| input.Ds10n.2L.wt.aro1.mi.pca.cmi.aracne.CLR.lite.bic.mf14.p10.json | output20190103163043 |
| input.Ds50n.2L.wt.aro1.mi.pca.cmi.aracne.CLR.lite.bic.mf14.json | output20180816154853 |
| input.Ds50n.2L.wt.aro1.mi.pca.cmi.aracne.CLR.lite.bic.mf14.p10.json | output20190103164415 |
| input.Ds100n.2L.wt.aro1.mi.pca.cmi.aracne.CLR.lite.bic.mf14.json | output20180816155654 |

Continued on next page

**Table 3.1 – continued from previous page**

| Input JSON File | Output Directory |
|---|---|
| input.Ds100n.2L.wt.aro1.mi.pca.cmi.aracne.CLR.lite.bic.mf14.p10.json | output20181218145844 |
| input.Ds100n.2L.wt.aro1.mi.pca.cmi.aracne.CLR.lite.bic.mf14.p3.json | output20190103170305 |
| input.Ds100n.2L.wt.aro1.mi.pca.cmi.aracne.CLR.lite.bic.mf14.p7.json | output20190103165930 |
| input.Ds100n.2L.wt.aro1.mi.pca.cmi.aracne.CLR.lite.bic.mf15.p10.json | output20181218152733 |
| input.Ds100n.2L.wt.aro1.mi.pca.cmi.aracne.CLR.lite.bic.mf16.p10.json | output20181219150849 |
| input.Ds100n.2L.wt.aro1.mi.pca.cmi.aracne.CLR.lite.bic.mf17.p10.json | output20181219153439 |
| input.Ds100n.2L.wt.aro1.mi.pca.cmi.aracne.CLR.lite.bic.mf18.p10.json | output20181218163804 |
| input.DmLc3E.aro1.mi.pca.cmi.CLR.lite.bic.mf15.p10.json | TGS-Lite-DmLc3E-output20180828124106 * |
| input.DmLc3L.aro1.mi.pca.cmi.CLR.lite.bic.mf15.p10.json | TGS-Lite-DmLc3L-output20180901132527 * |
| input.DmLc3P.aro1.mi.pca.cmi.CLR.lite.bic.mf15.p10.json | TGS-Lite-DmLc3P-output20180902203423 * |
| input.DmLc3A.aro1.mi.pca.cmi.CLR.lite.bic.mf15.p10.json | TGS-Lite-DmLc3A-output20180903010110 * |
| input.DmLc3E.aro1.mi.pca.cmi.aracne.CLR.lite.bic.mf15.p10.json | TGS-Lite-plus-DmLc3E-output20190123153500 * |
| input.DmLc3L.aro1.mi.pca.cmi.aracne.CLR.lite.bic.mf15.p10.json | TGS-Lite-plus-DmLc3L-output20190123161214 * |
| input.DmLc3P.aro1.mi.pca.cmi.aracne.CLR.lite.bic.mf15.p10.json | TGS-Lite-plus-DmLc3P-output20190123171659 * |
| input.DmLc3A.aro1.mi.pca.cmi.aracne.CLR.lite.bic.mf15.p10.json | TGS-Lite-plus-DmLc3A-output20190123181327 * |

### Chapter 4

### Appendix

#### 4.1 Memory Requirement of the Bene Step in *TGS*

The Bene step in *TGS* is based on the *Bene* algorithm [9]. This step takes a regulatee gene $v_j\_t_{(P+1)}$, its candidate regulator set $\mathcal{V}_{(j;p+1)}$ and their combined expression data $\mathcal{D}_{(\{v_j\_t_{(p+1)}\}\cup\mathcal{V}_{(j;(p+1))};\{t_p,t_{(p+1)}\};\mathcal{S})}$ as input. After analysing the data, it returns the highest scoring (w.r.t. the BIC scoring function) Bayesian network (BN) structure [9] where genes in $\mathcal{V}_{(j;p+1)}$ can only have outgoing edges and gene $v_j\_t_{(P+1)}$ can only have incoming edges. In order to find the highest scoring BN structure, the Bene step utilizes the following property: a BN is topologically a Directed Acyclic Graph (DAG) and a DAG is a directed graph with at least one topological order (a sequence of the nodes where parents of every node must precede the node in that sequence) [9]. Using this property, first the Bene step finds the best topological ordering among the given genes. For illustration, suppose that the candidate regulator set contains two candidate genes i.e. $\mathcal{V}_{(j;p+1)} = \{v_{i_1}\_t_{(p+1)}, v_{i_2}\_t_{(p+1)}\}$. Then either $\big(v_{i_1}\_t_{(p+1)}, v_{i_2}\_t_{(P+1)}, v_j\_t_{(p+1)}\big)$ or $\big(v_{i_2}\_t_{(p+1)}, v_{i_1}\_t_{(P+1)}, v_j\_t_{(p+1)}\big)$ can be the best topological order since only $v_j\_t_{(p+1)}$ can have incoming edges. Assume that the Bene step chooses $\big(v_{i_1}\_t_{(p+1)}, v_{i_2}\_t_{(p+1)}, v_j\_t_{(p+1)}\big)$ to be the best topological order. Given this order, it attempts to find the best set of parents for every node from the set of nodes preceding the former node in the provided order i.e. to find the best parents of $v_j\_t_{(p+1)}$ from $\{v_{i_1}\_t_{(p+1)}, v_{i_2}\_t_{(p+1)}\}$, to find the best parents of $v_{i_2}\_t_{(p+1)}$ from $\{v_{i_1}\_t_{(p+1)}\}$, and to find the best parents of $v_{i_1}\_t_{(p+1)}$ from $\emptyset$. However, the task boils down to finding the best parents of $v_j\_t_{(p+1)}$ from $\{v_{i_1}\_t_{(p+1)}, v_{i_2}\_t_{(p+1)}\}$ since $v_{i_1}\_t_{(p+1)}$ and $v_{i_2}\_t_{(p+1)}$ are not allowed to have parents (incoming edges). To accomplish this task, the Bene step creates two arrays, each of length $|\{v_{i_1}\_t_{(p+1)}, v_{i_2}\_t_{(p+1)}\}|$, where the $i^{th}$ element of the first array represents the $i^{th}$ subset of $\{v_{i_1}\_t_{(p+1)}, v_{i_2}\_t_{(p+1)}\}$ and the $i^{th}$ element of the second array stores the BIC score for the $i^{th}$ element of the first array (Figure 4.1). Subsequently, the subset with the highest BIC score is identified and the nodes in that subset are chosen as the parents of $v_j\_t_{(p+1)}$. Thus, the Bene step requires $\{v_{i_1}\_t_{(p+1)}, v_{i_2}\_t_{(p+1)}\} = |\mathcal{V}_{(j;p+1)}|$ subsets and that many scores to be held in memory.

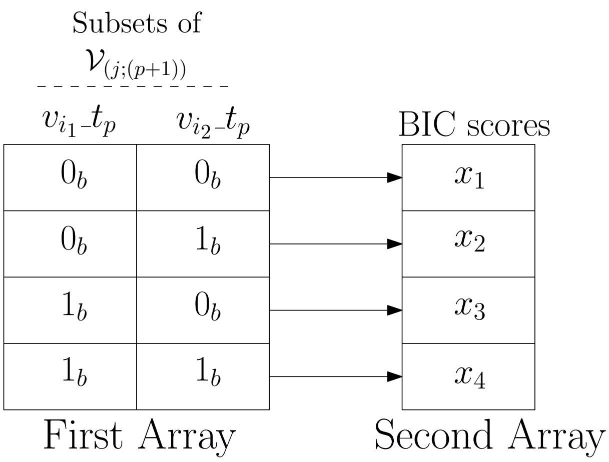

Figure 4.1: Memory Requirement of the Bene Step in the *TGS* Algorithm. The Bene step requires two arrays to be held in memory. The first array stores the subsets of a given candidate parent set, which is $\mathcal{V}_{(j;p+1)} = \{v_{i_1}\_t_{(p+1)}, v_{i_2}\_t_{(p+1)}\}$ in this example. Therefore, its subsets are $\{\emptyset, \{v_{i_2}\_t_p\}, \{v_{i_1}\_t_p\}, \{v_{i_1}\_t_p, v_{i_2}\_t_p\}\}$. Each subset is represented as a binary string of Boolean TRUE ($1_b$) and FALSE ($0_b$) values, indicating whether a node is present or not in that subset, respectively. The second array stores the BIC scores of these subsets. More specifically, the $i^{th}$ element of the second array contains the BIC score of the BN structure where the genes in the $i^{th}$ element of the first array are the parents of a given target node, which is $v_j\_t_{(p+1)}$ in this example.

#### 4.2 Time comlexity of BIC Score Computation

Computation of the BIC score of a given candidate regulator set for a given target gene during a particular time interval is performed using the following C function.

```
/* ============================================================================
* The following function, namely 'log_likelihood()',
* is a part of the C program file 'score_bn.c'.
* URL: https://github.com/sap01/TGS-Lite/blob/master/src/score_bn.c
* Words 'node' and 'gene' are used interchangeably.
* Words 'parent' and 'regulator' are used interchangeably.
* ============================================================================ */
/*
* Input parameters:
* d = data.
* n_nodes = number of nodes.
* n_cases = number of replicates per node per time point.
* ns = an array containing node sizes i.e. the discrete levels of nodes;
*         ns[i] = number of discrete levls of the i-th node.
* ni = node index of the target node.
* pa = an array containing the indices of the candidate parent nodes.
* n_pa = number of candidate parent nodes.
* penalty = sparsity penalty to be deducted from the computed
*         log likelihood for the given scoing function.
*/
double log_likelihood ( unsigned int * d, unsigned int n_nodes, unsigned int n_cases,
unsigned int * ns, unsigned int ni, unsigned int * pa, unsigned int n_pa,
double penalty ) {
        int i, j, index, elmt, stride;
        double acc, logl;
```

```c
int cum_prod_sizes[n_pa+2];
cum_prod_sizes[0] = 1;
cum_prod_sizes[1] = ns[ni];

/*
* Time complexity of the following for loop
* = n_pa
* = number of candidate parents of the given target node
* = O(M_f).
*/
for( i = 0; i < n_pa; i++ )
        cum_prod_sizes[i+2] = cum_prod_sizes[i+1] * ns[pa[i]];

int strides[n_pa + 1];
strides[0] = ni * n_cases;

/*
* Time complexity of the following for loop
* = n_pa
* = number of candidate parents of the given target node
* = O(M_f).
*/
for( i = 0; i < n_pa; i++ )
        strides[i+1] = pa[i] * n_cases;

int prod_sizes = cum_prod_sizes[n_pa+1];
int prod_sizes_pa = prod_sizes / ns[ni];

double * counts = calloc( prod_sizes, sizeof(double) );

/*
* Number of iterations for the following for loop
* = n_cases
* = S.
*/
for( i = 0; i < n_cases; i++ ) {
        index = 0;

        /*
        * Time complexity of the following for loop
        * = (n_pa + 1)
        * = (number of candidate parents of the given target node + 1)
        * = (O(M_f) + 1).
        */
        for( j = 0; j < n_pa + 1; j++ ) {
                elmt = d[ i + strides[j] ];
                if( elmt == NA_INTEGER )
                break;

                index += (elmt - 1) * cum_prod_sizes[j];
        }

        if( j < n_pa + 1 )
```

```
                        continue;

                        counts[index] += 1;
        }

        logl = 0.0;

        /*
        * Number of iterations for the following for loop
        * = prod_sizes_pa
        * = product of the numbers of discrete levels of
        *          the candidate parent nodes.
        */
        for( i = 0; i < prod_sizes_pa; i++ )
        {
                stride = i * ns[ni];
                acc    = counts[stride];

                /*
                * Time complexity of the following for loop
                * = (ns[ni] - 1)
                * = (Number of discrete levels of the target node - 1).
                */
                for( j = 1; j < ns[ni]; j++ )
                {
                        acc  += counts[stride + j];
                }

                acc = log(acc + 1);

                /*
                * Time complexity of the following for loop
                * = (ns[ni] - 1)
                * = (Number of discrete levels of the target node - 1).
                */
                for ( j = 1 ; j < ns[ni] ; j++ )
                {
                        logl += counts[stride + j] * (log(counts[stride + j] + 1)
                         - acc);
                }

        }

        logl -= penalty * prod_sizes_pa * (ns[ni] - 1);

        free(counts);

        return logl;
}
```

From the above C function , the time complexity of the BIC score computation can be derived as follows:

$$T_{\text{BIC}}\left(V;T;S;M_f;\vec{\delta}\right) = \mathcal{O}\left(M_f\right) + \mathcal{O}\left(M_f\right) + S\left(\mathcal{O}\left(M_f\right)+1\right) + \left(\prod_{v_i\_t_p \in curr.set} \delta_i\right) \times 2 \times \left(\delta_j - 1\right)$$

(where 'curr.set' = the given candidate regulator set;

$v_j\_t_{(p+1)}$ = the given target/regulatee gene;

$v_i\_t_p$ = an arbitrary candidate regulator gene $\in$ curr.set;

$\delta_i$ = number of discrete levels of gene $v_i\_t_p$ = number of discrete levels of gene $v_i$;

$\delta_j$ = number of discrete levels of gene $v_j\_t_{(p+1)}$ = number of discrete levels of gene $v_j$;

$\vec{\delta}$ = a vector containing the numbers of discrete levels of genes, e.g., $i^{\text{th}}$ element of $\vec{\delta}$ = number of discrete levels of gene $v_i = \delta_i$.)

$$= \mathcal{O}\left(M_f\right) + S\left(\mathcal{O}\left(M_f\right)+1\right) + \left(\prod_{v_i\_t_p \in curr.set} \delta_i\right) \times 2 \times \left(\delta_j - 1\right)$$

$(\because \mathcal{O}\left(M_f\right) + \mathcal{O}\left(M_f\right) = \mathcal{O}\left(M_f\right))$

$$= \mathcal{O}\left(M_f\right) + S \times \mathcal{O}\left(M_f\right) + S + \left(\prod_{v_i\_t_p \in curr.set} \delta_i\right) \times 2 \times \left(\delta_j - 1\right)$$

$$= (1+S) \times \mathcal{O}\left(M_f\right) + S + \left(\prod_{v_i\_t_p \in curr.set} \delta_i\right) \times 2 \times \left(\delta_j - 1\right). \quad (4.1)$$

From Equation 4.1 , the time complexity can be transformed into an expression only w.r.t. $V$ and $M_f$ as follows:

$$T_{\text{BIC}}\left(V;M_f\right) = (1+\Theta(1)) \times \mathcal{O}\left(M_f\right) + \Theta(1) + \left(\prod_{v_i\_t_p \in curr.set} \delta_i\right) \times 2 \times \left(\delta_j - 1\right)$$

($\because$ $S$ is independent of $V$ and $M_f$ i.e. $S = \Theta(1)$)

$$= \Theta(1) \times \mathcal{O}\left(M_f\right) + \Theta(1) + \left(\prod_{v_i\_t_p \in curr.set} \delta_i\right) \times 2 \times \left(\delta_j - 1\right)$$

$(\because (1+\Theta(1)) = \Theta(1))$

$$= \mathcal{O}\left(M_f\right) + \Theta(1) + \left(\prod_{v_i\_t_p \in curr.set} \delta_i\right) \times 2 \times \left(\delta_j - 1\right) \quad (4.2)$$

$(\because (\Theta(1) \times \mathcal{O}\left(M_f\right)) = \mathcal{O}\left(M_f\right)))$.

$$= \mathcal{O}\left(M_f\right) + \Theta(1) + \left(\prod_{v_i\_t_p \in curr.set} \Theta(1)\right) \times 2 \times \left(\Theta(1) - 1\right)$$

($\because$ $\delta_i, \delta_j$ are independent of $V$ and $M_f$ for all $\delta_i, \delta_j \in \vec{\delta}$)

$$= \mathcal{O}\left(M_f\right) + \Theta(1) + \Theta(1)$$
$$= \mathcal{O}\left(M_f\right) + \Theta(1). \quad (4.3)$$

#### 4.3 Time comlexity of ***Find-best-set-Lite***

The time complexity of Algorithm *Find-best-set-Lite* (Algorithm TODO of the main paper) can be derived as follows:

$$
\begin{aligned}
&T_{\text{Find-best-set-Lite}}\left(V;T;S;M_f;\vec{\delta}\right)\\
&= T_{\text{BIC}}\left(V;T;S;M_f;\vec{\delta}\right) + \mathcal{O}\big(2^{M_f}-1\big) \times \left(\mathcal{O}(M_f) + T_{\text{BIC}}\left(V;T;S;M_f;\vec{\delta}\right)\right).
\end{aligned} \tag{4.4}
$$

#### 4.4 Time comlexity of ***TGS-Lite***

The time complexity of Algorithm *TGS-Lite* (Algorithm TODO of the main paper) can be derived as follows:

$$
\begin{aligned}
&T_{\text{TGS-Lite}}\left(V;T;S;M_f;\vec{\delta}\right)\\
&= \mathcal{O}\left(V^2\right) + \mathcal{O}\left(V^2\right) + V \times (T-1) \times \left(\mathcal{O}\left(V\right) + \mathcal{O}\left(M_f\right) + T_{\text{Find-best-set-Lite}}\left(V;T;S;M_f;\vec{\delta}\right) + \mathcal{O}(M_f)\right)\\
&= \mathcal{O}\left(V^2\right) + \mathcal{O}\left(V^2\right) + V \times (T-1) \times \Bigg(\mathcal{O}\left(V\right) + \mathcal{O}\left(M_f\right) + T_{\text{BIC}}\left(V;T;S;M_f;\vec{\delta}\right)\\
&\qquad + \mathcal{O}\big(2^{M_f}-1\big) \times \left(\mathcal{O}(M_f) + T_{\text{BIC}}\left(V;T;S;M_f;\vec{\delta}\right)\right) + \mathcal{O}(M_f)\Bigg)\\
&\qquad \text{(From Equation 4.4 )}\\
&= \mathcal{O}\left(V^2\right) + V \times (T-1) \times \Bigg(\mathcal{O}\left(V\right) + \mathcal{O}\left(M_f\right) + T_{\text{BIC}}\left(V;T;S;M_f;\vec{\delta}\right)\\
&\qquad + \mathcal{O}\big(2^{M_f}-1\big) \times \left(\mathcal{O}(M_f) + T_{\text{BIC}}\left(V;T;S;M_f;\vec{\delta}\right)\right)\Bigg).
\end{aligned} \tag{4.5}
$$

$$\left(\because \mathcal{O}\left(V^2\right) + \mathcal{O}\left(V^2\right) = \mathcal{O}\left(V^2\right);\ \mathcal{O}\left(M_f\right) + \mathcal{O}\left(M_f\right) = \mathcal{O}\left(M_f\right).\right)$$

From Equation 4.5 , the time complexity can be transformed into an expression only w.r.t. $V$ and $M_f$ as follows:

$$
\begin{aligned}
&T_{\text{TGS-Lite}}\left(V; M_f\right)\\
&= \mathcal{O}\left(V^2\right) + V \times (T-1) \times \Bigg(\mathcal{O}\left(V\right) + \mathcal{O}\left(M_f\right) + \Big(\mathcal{O}(M_f) + \Theta(1)\Big)\\
&\qquad + \mathcal{O}\big(2^{M_f}-1\big) \times \Big(\mathcal{O}(M_f) + \Big(\mathcal{O}(M_f) + \Theta(1)\Big)\Big)\Bigg)\\
&\qquad \text{(From Equation 4.3 )}\\
&= \mathcal{O}\left(V^2\right) + V \times (T-1) \times \Bigg(\mathcal{O}\left(V\right) + \mathcal{O}\left(M_f\right) + \mathcal{O}(M_f) + \Theta(1)\\
&\qquad + \mathcal{O}\big(2^{M_f}-1\big) \times \Big(\mathcal{O}(M_f) + \mathcal{O}(M_f) + \Theta(1)\Big)\Bigg)\\
&= \mathcal{O}\left(V^2\right) + V \times (T-1) \times \Bigg(\mathcal{O}\left(V\right) + \mathcal{O}\left(M_f\right) + \Theta(1) + \mathcal{O}\big(2^{M_f}-1\big) \times \Big(\mathcal{O}(M_f) + \Theta(1)\Big)\Bigg)\\
&\qquad (\because \mathcal{O}\left(M_f\right) + \mathcal{O}\left(M_f\right) = \mathcal{O}\left(M_f\right))\\
&= \mathcal{O}\left(V^2\right) + V \times \Theta\left(1\right) \times \Bigg(\mathcal{O}\left(V\right) + \mathcal{O}\left(M_f\right) + \Theta(1) + \mathcal{O}\big(2^{M_f}-1\big) \times \Big(\mathcal{O}(M_f) + \Theta(1)\Big)\Bigg)\\
&\qquad \Big(\because T \text{ is independent of } V \text{ and } M_f \text{ i.e. } (T-1) = \Theta(1)\Big)\\
&= \mathcal{O}\left(V^2\right) + \Theta\left(V\right) \times \Bigg(\mathcal{O}\left(V\right) + \mathcal{O}\left(M_f\right) + \Theta(1) + \mathcal{O}\big(2^{M_f}-1\big) \times \Big(\mathcal{O}(M_f) + \Theta(1)\Big)\Bigg). \qquad (4.6)\\
&\qquad \because \Big(V \times \Theta\left(1\right) = \Theta\left(V\right).\Big)
\end{aligned}
$$

Furthermore, it can be noted that $M_f$ is not independent of $V$. The value of $M_f$ is upper bounded by that of $V$; therefore in theory, $M_f \leq V$. However in practice, $M_f$ is expected to be much smaller than $V$ ($M_f \ll V$); hence, Pyne et al. [7] considers that:

$$M_f = o(\lg V) \tag{4.7}$$

Using this relationship between $M_f$ and $V$, Equation 4.6 can be transformed further into an expression only w.r.t. $V$ as follows:

$$
\begin{aligned}
&T_{\text{TGS-Lite}}(V) \\
&= \mathcal{O}\left(V^2\right) + \Theta(V) \times \left(\mathcal{O}(V) + o(\lg V) + \Theta(1) + \mathcal{O}\Big(2^{o(\lg V)} - 1\Big) \times \Big(o(\lg V) + \Theta(1)\Big)\right) \\
&\qquad \because \Big(M_f = o(\lg V) \implies \mathcal{O}(M_f) = \mathcal{O}(o(\lg V)) = o(\lg V).\Big) \\
&= \mathcal{O}\left(V^2\right) + \Theta(V) \times \left(\mathcal{O}(V) + o(\lg V) + \Theta(1) + \mathcal{O}(o(V) - 1) \times \Big(o(\lg V) + \Theta(1)\Big)\right) \\
&\qquad \because \Big(2^{o(\lg V)} = o(V).\Big) \\
&= \mathcal{O}\left(V^2\right) + \Theta(V) \times \left(\mathcal{O}(V) + o(\lg V) + \Theta(1) + \mathcal{O}(o(V)) \times \Big(o(\lg V) + \Theta(1)\Big)\right) \\
&\qquad \because \Big(o(V) - 1 = o(V).\Big) \\
&= \mathcal{O}\left(V^2\right) + \Theta(V) \times \left(\mathcal{O}(V) + o(\lg V) + \Theta(1) + o(V) \times \Big(o(\lg V) + \Theta(1)\Big)\right) \\
&\qquad \because \Big(\mathcal{O}(o(V)) = o(V).\Big) \\
&= \mathcal{O}\left(V^2\right) + \Theta(V) \times \left(\mathcal{O}(V) + o(\lg V) + \Theta(1) + o(V) \times o(\lg V) + o(V) \times \Theta(1)\right) \\
&= \mathcal{O}\left(V^2\right) + \Theta(V) \times \left(\mathcal{O}(V) + o(\lg V) + \Theta(1) + o(V \lg V) + o(V)\right) \\
&= \mathcal{O}\left(V^2\right) + \Theta(V) \times \left(o(V \lg V) + \Theta(1)\right) \\
&\qquad \because \Big(\mathcal{O}(V) + o(\lg V) + o(V \lg V) + o(V) = o(V \lg V).\Big) \\
&= \mathcal{O}\left(V^2\right) + \Theta(V) \times o(V \lg V) + \Theta(V) \times \Theta(1) \\
&= \mathcal{O}\left(V^2\right) + o\big(V^2 \lg V\big) + \Theta(V) \times \Theta(1) \\
&\qquad \because \Big(\Theta(V) \times o(V \lg V) = o\big(V^2 \lg V\big).\Big) \\
&= \mathcal{O}\left(V^2\right) + o\big(V^2 \lg V\big) + \Theta(V) \\
&\qquad \because \Big(\Theta(V) \times \Theta(1) = \Theta(V).\Big) \\
&= o\big(V^2 \ \lg V\big).
\end{aligned} \tag{4.8}
$$

#### 4.5 Time comlexity of *TGS-Lite+*

The time complexity of the *TGS-Lite+* algorithm (Algorithm TODO of the main paper) w.r.t. $V$ can be derived as follows:

$$
\begin{aligned}
&T_{\text{TGS-Lite+}}(V) \\
&= \mathcal{O}\big(V^3\big) + T_{\text{TGS-Lite}}(V) \\
&= \mathcal{O}\big(V^3\big) + o\big(V^2 \ \lg V\big) && \text{(From Equation 4.8 )} \\
&= \mathcal{O}\big(V^3\big).
\end{aligned} \tag{4.9}
$$

#### 4.6 Time comlexity of *TGS*

The time complexity of algorithm *TGS* with the max fan-in restriction (Algorithm 4 of Pyne et al. [7] ) can be derived as follows:

$$\begin{aligned} & T_{\text{TGS}}\,(V) \\ & = \mathcal{O}\big(V^2\big) + o\Big(\,(T-1) \times V \times {M_f}^2 \times 2^{\left(M_f - 2\right)}\Big). \\ & \qquad \because \text{(From Equation 1 of Pyne et al. [7] )} \end{aligned} \tag{4.10}$$

In the RHS of this equation, the sub-expression $o\Big({M_f}^2 \times 2^{\left(M_f-2\right)}\Big)$ is used by Pyne et al. to represent the time complexity of the *Bene* step. This sub-expression is derived directly by substituting $n$ with $M_f$ in the time complexity of the original *Bene* algorithm, which is $o\big(n^2 2^{(n-2)}\big)$ where $n$ = total number of genes processed by *Bene* [9]. While the sub-expression is correct, it can be reduced to a tighter bound since the *Bene* step in *TGS* has to perform a strictly smaller number of tasks than that of the original *Bene* algorithm as explained below.

The original *Bene* algorithm attempts to find the best candidate regulator set for every given gene. It assumes that every gene is a candidate regulator of every other gene. The original *Bene* algorithm requires $o\big((n-1)\,2^{(n-2)}\big)$ time for finding the best parents (the best candidate regulator set) for every gene (Section 3.2 of Silander et al. [9]). Therefore to find the best candidate regulator sets for all $n$ genes, it requires a time complexity of $o\big(n\,(n-1)\,2^{(n-2)}\big) = o\big(n^2 2^{(n-2)}\big)$.

On the other hand, the *Bene* step in the *TGS* algorithm attempts to find the best candidate regulator set of a single gene, pre-specified as the regulatee gene; all other genes are considered as the former gene's candidate regulators. Therefore, the time required by the *Bene* step for finding the best candidate regulator set of the pre-specified regulatee gene is $o\big((n-1)\,2^{(n-2)}\big)$. Now, for the *Bene* step:

$$\begin{aligned} n &= \text{number of regulatee genes} + \text{number of candidate regulator genes} \\ &= 1 + \mathcal{O}(M_f). \end{aligned} \tag{4.11}$$

Hence, time complexity of the *Bene* step:

$$\begin{aligned} T_{\text{Bene step}}\,(V; M_f) &= o\Big(\,(n-1)\,2^{(n-2)}\Big) \\ &= o\Big(\,(1+\mathcal{O}(M_f)-1)\,2^{\left(1+\mathcal{O}(M_f)-2\right)}\Big) \\ &\qquad \text{(From Equation 4.11 )} \\ &= o\Big(\mathcal{O}(M_f)\ 2^{\left(\mathcal{O}(M_f)-1\right)}\Big) \\ &= o\Big(\mathcal{O}(M_f)\ 2^{\left(o(M_f)\right)}\Big) \\ &= o\big(M_f\ 2^{M_f}\big). \end{aligned} \tag{4.12}$$

Therefore, the time complexity of algorithm *TGS* with the max fan-in restriction can be revised as follows:

$$
\begin{aligned}
&T_{\text{TGS}}(V) \\
&= \mathcal{O}\big(V^2\big) + o\Big(\,(T-1) \times V \times M_f \times 2^{M_f}\Big) \\
&\qquad\text{(Substituting sub-expression } \Big({M_f}^2 \times 2^{\big(M_f - 2\big)}\Big) \text{ in Equation 4.10 with the RHS in Equation 4.12 )} \\
&= \mathcal{O}\big(V^2\big) + o\Big(\Theta(1) \times V \times M_f \times 2^{M_f}\Big) \\
&\qquad \because (T \text{ is independent of } V \text{ i.e. } (T-1) = \Theta(1).) \\
&= \mathcal{O}\big(V^2\big) + o\Big(\Theta(V) \times M_f \times 2^{M_f}\Big) \\
&\qquad \because \Big(\Theta(1) \times V = \Theta(V).\Big) \\
&= \mathcal{O}\big(V^2\big) + o\Big(\Theta(V) \times o(\lg V) \times 2^{(o(\lg V))}\Big) \\
&\qquad \text{(From Equation 4.7 )} \\
&= \mathcal{O}\big(V^2\big) + o\Big(\Theta(V) \times o(\lg V) \times o(V)\Big) \\
&\qquad \because \Big(2^{(o(\lg V))} = o(V).\Big) \\
&= \mathcal{O}\big(V^2\big) + o\Big(o(V \times \lg V \times V)\Big) \\
&= \mathcal{O}\big(V^2\big) + o\Big(o\big(V^2 \times \lg V\big)\Big) \\
&= \mathcal{O}\big(V^2\big) + o\Big(V^2 \ \lg V\Big) \\
&= o\Big(V^2 \ \lg V\Big).
\end{aligned} \tag{4.13}
$$

#### 4.7 Time comlexity of *TGS+*

The time complexity of the *TGS+* algorithm with the max fan-in restriction (Algorithm 6 of Pyne et al. [7] ) can be derived w.r.t. $V$ as follows:

$$
\begin{aligned}
&T_{\text{TGS+}}(V) \\
&= \mathcal{O}\big(V^3\big) + T_{\text{TGS}}(V) \\
&= \mathcal{O}\big(V^3\big) + o\big(V^2 \ \lg V\big) \qquad \text{(From Equation 4.13 )} \\
&= \mathcal{O}\big(V^3\big).
\end{aligned} \tag{4.14}
$$

#### 4.8 Evaluation Metrics for Comparing a Predicted Network with the True Network

Given a dataset and the corresponding true network that generated the data, the following metrics are used to evaluate the learning power of an algorithm. True Positive (TP) and False Positive (FP) stand for number of edges correctly predicted and number of edges incorrectly predicted, respectively. On the other hand, True Negative (TN) and False Negative (FN) represent number of non-edges (absence of edges in the true network) correctly predicted and number of non-edges incorrectly predicted, respectively.

- True Positive Rate (TPR): (Synonyms: Sensitivity, Recall)
  $TPR = TP/(TP + FN)$.
- False Positive Rate (FPR):
  $FPR = FP/(FP + TN)$.
- False Discovery Rate (FDR):
  $FDR = FP/(TP + FP)$.
- Positive Predictive Value (PPV): (Synonym: Precision)
  $PPV = TP/(TP + FP)$.
- Overall Accuracy(ACC):
  $ACC = (TP + TN)/(TP + TN + FP + FN)$.
- Matthew's correlation coefficient (MCC):
  $MCC = \frac{TP \times TN - FP \times FN}{\sqrt{(TP+FP)(TP+FN)(TN+FP)(TN+FN)}}$ .
- F1-Score:
  $F1 = (2 \times PPV \times TPR)/(PPV + TPR)$. When $PPV$ and $TPR$ are zeros, $F1$ is considered as zero [1].

#### 4.9 A Comparative Study Against the Performance of the Winning Team in DREAM3 In Silico Network Challenge

In this section, a comparative study is presented between the performances of the proposed algorithms and those of the winning team's algorithm in DREAM3 In Silico Network Challenge [6]. The reason behind conducting this comparison is that the in-silico benchmark datasets used to evaluate the performance of the proposed algorithms (in Section 5.1 of the main paper) are a subset of the datasets used in the aforementioned challenge. Therefore, it is interesting to find out how the proposed algorithms perform compared to the winning team's algorithm on the same datasets.

The datasets in question are called Ds10n, Ds50n and Ds100n. Dataset Ds10n corresponds to the time-series dataset generated with the 'Yeast1' network under the 10-node sub-challenge of the DREAM3 challenge. Similarly, datasets Ds50n and Ds100n correspond to the time-series datasets generated with the 'Yeast1' networks under the 50-node and 100-node sub-challenges, respectively, of the DREAM3 challenge.

The winning team, also known as the 'B Team', reports the performances of their algorithm (henceforth, *'BTA'*, an abbreviation for 'B Team's Algorithm') in Yip et al. [10]. For the 10-node sub-challenge with 'Yeast1' network, *BTA* achieves an AUROC value of 0.949. However, it uses three datasets: (1) dataset Ds10n, (2) a knock-out dataset where each gene's expression is knocked out to zero, one gene at a time, while other genes' expressions are recorded, and (3) a knock-down dataset where each gene's expression is knocked down to half of its normal ('wild-type') expression, one gene at a time, while other genes' expressions are recorded. Yip et al. demonstrates that the knock-out dataset is especially crucial to *BTA* for identifying edges where a gene has a single regulator. On the other hand, the proposed algorithms, namely *TGS-Lite* and *TGS-Lite+*, only use dataset Ds10n.

It is not feasible to find out AUROC values for *TGS-Lite* and *TGS-Lite+* that will be comparable to the aforementioned AUROC value of *BTA*. The reason lies in the difference between the outputs of these algorithms. *BTA* outputs a ranked list of all possible edges but not a network. The rank of an edge depends upon a real number which represents *BTA*'s confidence on the occurrence of that edge. The edges are listed in the decreasing order of their confidence values. The list contains $V(V-1)$ edges, where $V =$ number of genes, since self loops are not allowed. Therefore, generating a network from the list requires a user-defined parameter '$k$', which selects the top $k$ edges from the list. Thus, the aforementioned AUROC value is generated by varying the value of $k$ from 1 to $V(V-1)$. On the other hand, both *TGS-Lite* and *TGS-Lite+* directly output networks without the need of any user-defined parameters. Hence, it is not feasible to generate their ROC curves equivalent to that of *BTA*.

However, it might be possible to make an indirect comparison between *BTA* and the proposed algorithms. That requires an in-depth look into the way *BTA* creates the ranked list. It generates the list through seven steps, called 'batch 1' to 'batch 7'. During batches 1 to 6, *BTA* makes a ranked sub-list of potentially true edges along with their confidence scores. The remaining edges are then appended to the sub-list; this step is known as batch 7. Therefore, batch 7 edges do not alter the ranks of edges in the initial sub-list. The experimental results show that the initial sub-list contains the strongest predictions whereas batch 7 contains the weakest ones [10]. For example, in case of the 'size 10 network' of 'Yeast1', the initial sub-list contains 32 edges out of which 9 are correct predictions; on the contrary, batch 7 predicts a total of 58 edges out of which only 1 turns out to be correct. Therefore, only the initial sub-list is considered to generate the predicted network, thus minimizing the weakness of *BTA*. This network is then compared against the rolled networks of *TGS-Lite* and *TGS-Lite+*.

Following the aforementioned comparative strategy, it is observed that *BTA* predicts a higher number of true positive edges. However, it does so at the cost of a significantly higher number of false positive ones. *BTA* predicts a total of 32 edges out of which only 9 are true positives. On the other hand, *TGS-Lite* predicts 13 edges out of which 3 are true positives, and *TGS-Lite+* predicts only 4 edges out of which 3 turn out to be true positives. Therefore, *TGS-Lite+* achieves the best balance between the true positives and false positives. The same pattern is observed for the 50-node and 100-node datasets (Table 4.1). These results provide further support to *TGS-Lite+*'s superior reconstruction power.

Moreover, Yip et al. discusses that the runtime is a major limitation of *BTA*. For the 100-node dataset, it takes 78 hours on a 'high-end cluster' (cluster configuration is not found) [10]. In contrast,

Table 4.1: A Comparison between Algorithm *BTA* and the Proposed Algorithms on the chosen DREAM3 Datasets. Here, 'x / y' represents the 'true positives / false positives' predicted by the corresponding algortihm (column) for a given dataset (row). Please note that *BTA* uses time-series, knock-out and knock-down data of a given dataset; on the other hand, *TGS-Lite* and *TGS-Lite+* use only the time-series data.

| **Dataset** | ***BTA*** | ***TGS-Lite*** | ***TGS-Lite+*** |
|---|---|---|---|
| 10-node | 9/23 | 3/10 | 3/1 |
| 50-node | 69/586 | 15/342 | 6/100 |
| 100-node | 145/2187 | 28/790 | 19/181 |

*TGS-Lite+* takes around 12 minutes on 3.07GHz CPU with 32 GB main memory. However, it must be noted that *BTA* processes time-series, knock-out and knock-down data, while *TGS-Lite+* processes only the time-series data.

**Summary:** The results of this comparative study can be summarised as follows:

- *BTA* outputs a ranked list of all possible edges, not a network. Even when a some strategy is employed to generate a network from the ranked list, the resulting network is a summary GRN. Therefore, the time interval(s) during which an edge is active can not be known. Thus, *BTA* does not address the problem of time-varying GRN reconstruction, which the proposed algorithms do.
- *BTA* predicts higher numbers of true positives than the proposed algorithms. However, it does so at the cost of incurring significantly higher numbers of false positives. *TGS-Lite+* provides a better balance in predictions of true and false positives.
- *BTA* requires much longer runtime than those of the proposed algorithms.
- The proposed algorithms only use time-series gene expression data. On the other hand, *BTA* can also utilise knock-out and knock-down gene expression data. Especially, knock-out data is proved to be very useful to *BTA* for identifying simple regulatory relationships, such as - when a gene has a single regulator [10]. Therefore, the reconstruction powers of the proposed algorithms might be further enhanced by adding the capability for utilising knock-out and knock-down gene expression data. It can be a worthwhile direction to explore in future.

#### 4.10 Explanation for the Retention of *TGS-Lite+*'s Recall with Ds10n

This section presents an explanation for the observation that *TGS-Lite+* achieves a significantly higher recall (0.75) than that of *TGS-Lite* (0.231) for dataset Ds10n, while retaining the same recall (0.3).

By design, *TGS-Lite+* is expected to achieve a significantly higher precision than that of *TGS-Lite* while sacrificing a reasonable amount of recall, if necessary. For datasets Ds50n and Ds100n, *TGS-Lite+* does achieve significantly higher precisions at the cost of slightly lower recalls. However, for dataset Ds10n, *TGS-Lite+* is able to achieve higher precision without making any compromise on recall.

The reason behind *TGS-Lite+*'s aforementioned achievement is due to its difference with *TGS-Lite* and how that difference is favoured by the underlying true network of Ds10n. The difference is that *TGS-Lite+* has an additional step compared to *TGS-Lite*; it is known as the ARACNE step. This step is used to refine the raw mutual information matrix calculated from the given data. While *TGS-Lite* uses the raw mutual information matrix to shortlist candidate regulators, *TGS-Lite+* uses the refined mutual information matrix to do so.

The refined mutual information matrix is produced through the ARACNE step by eliminating positive ($\in \mathbb{R}^+$) mutual information values which are potentially due to indirect regulatory relationships. This step assumes that genes $v_1$ and $v_2$ does not have a direct regulatory relationship if there exists at least another gene $v_3$ such that the mutual information between the former genes is smaller than their mutual information with $v_3$; formally, $\mathscr{M}(v_1, v_2) < \min(\mathscr{M}(v_1, v_3), \mathscr{M}(v_3, v_2))$. If that is the case, then it is assumed that $v_1$ and $v_2$ have an indirect regulatory relationship via $v_3$. For example, $v_1$ regulates $v_3$ and $v_3$ in turn regulates $v_2$. In such a situation, the ARACNE step replaces the $\mathscr{M}(v_1, v_2)$ entry in the mutual information matrix with zero. Therefore, $v_1$ is not allowed to be a candidate regulator of $v_2$ and vice versa.

The aforementioned refinement strategy is proposed by Margolin et al. [5]. Based on this strategy, they design an algorithm, known as *ARACNE*, after which the concerned *TGS-Lite+* step is named. This strategy is demonstrated by Margolin et al. and Pyne et al. [7] to significantly reduce false positive edges at a reasonable loss of true positive edges. Pyne et al. discusses one of the cases where true positive edges are lost. This case happens when two genes $v_1$ and $v_2$ share a direct as well as an indirect regulatory relationships. For example, when $v_1$ regulates $v_2$ and another gene $v_3$ while $v_3$ also regulates $v_2$ (Figure 4.2). In that case, the edge from $v_1$ to $v_2$ is known as a feed-forward edge.

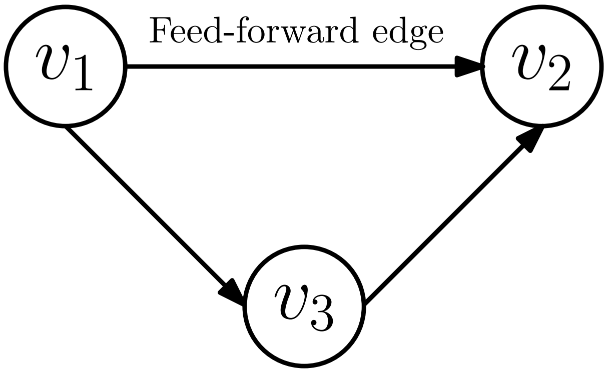

Figure 4.2: An example of a Feed-forward Edge. In this example, gene $v_1$ regulates genes $v_2$ and $v_3$. In addition, $v_3$ regulates $v_2$. Therefore, $v_1$ has a direct regulatory relationship with $v_2$ as well as an indirect regulatory relationship with $v_2$ via $v_3$. In this case, the edge from $v_1$ to $v_2$ is called a feed-forward edge. Such edges are a known characteristic of biological networks [4]. They are believed to improve robustness of the network, e.g. in the given situation, $v_2$ can be directly regulated through $v_1$ even when $v_3$ is not under control.

In the aforementioned example, the ARACNE step rejects the feed-forward edge if $\mathscr{M}(v_1, v_2) < \min(\mathscr{M}(v_1, v_3), \mathscr{M}(v_3, v_2))$. In that case, *TGS-Lite+* loses a true edge. Hence, the lower the number of feed-forward edges in the underlying network, the lower the chances of *TGS-Lite+* predicting a lower number of true positive edges than that of *TGS-Lite*. Therefore, it is expected that *TGS-Lite+* would exhibit a smaller loss in recall with Ds10n than with Ds50n and Ds100n, since only 10% edges of Ds10n's true network are feed-forward edges whereas such edges are around 39% of true edges in case of Ds50n and Ds100n [7].

However, it is interesting that *TGS-Lite+* exhibits absolutely no loss in recall with Ds10n. *TGS-Lite+*, like *TGS-Lite*, correctly predicts 3 out of 10 true edges. Among those 3 edges, only 2 are predicted by both the algorithms. Therefore, each algorithm correctly predicts a unique edge. The unique edge correctly predicted by *TGS-Lite* is the edge from gene G3 to gene G5. It is a feed-forward edge (Figure 4.3).

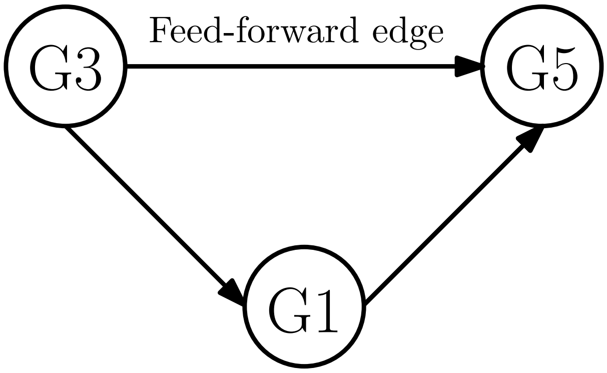

Figure 4.3: The Feed-forward edge in the True Network underlying Dataset Ds10n. Only a sub-network is shown. The complete network has ten genes, namely {G1, ..., G10}, and ten edges.

*TGS-Lite+* rejects this feed-forward edge. Ideally, the reason for the rejection should be the assumption that G3 regulates G5 only via G1. *TGS-Lite+* can make that assumption only if: $\mathscr{M}(G3, G5) < \min(\mathscr{M}(G3, G1), \mathscr{M}(G1, G5))$. However, this criterion does not satisfy. Instead, the rejection is caused by genes G4 and G6. *TGS-Lite+* assumes that G3 regulates G5 via G4 or G6 since the following criteria are satisfied:

- $\mathscr{M}(G3, G5) < \min(\mathscr{M}(G3, G4), \mathscr{M}(G4, G5))$,
- $\mathscr{M}(G3, G5) < \min(\mathscr{M}(G3, G6), \mathscr{M}(G6, G5))$.

The aforementioned criteria can be satisfied for multiple reasons. One of them is the inaccurate calculations of mutual information values. That can be caused by:

- noise in the data,
- information loss due to data discretization,
- not considering the temporal order in data during calculation, etc.

It remains an open challenge to devise new methods to calculate mutual information more accurately from noisy time-series data.

*TGS-Lite+*, on the other hand, uniquely predicts the true edge from G1 to G2. According to the raw mutual information matrix, G2 shares significantly higher mutual information with: G1, G4 and G9, compared with other genes. Therefore, these three genes should be shortlisted as G2's candidate regulators. However, the ARACNE step eliminates G4 from the shortlist, assuming that G4 regulates G2 via G1 since: $\mathscr{M}(G4, G2) < \min(\mathscr{M}(G4, G1), \mathscr{M}(G1, G2))$. Thus, the candidate regulator set of G2 becomes $\{G1, G9\}$. Among all its subsets, $\{G1\}$ achieves the highest BIC score ($= -2$). Hence, G1 gets selected as the sole regulator of G2.

In contrast, *TGS-Lite* has to use $\{G1, G4, G9\}$ as the candidate regulator set of G2. Among its subsets, $\{G1\}$ and $\{G4\}$ jointly achieve the highest BIC score ($= -2$). According to the current implementation of *TGS-Lite*, the tie is broken in favour of the higher indexed gene i.e. G4. Hence, G4 is selected as the sole regulator of G2, which is a false positive prediction.

Thus, in case of Ds10n, the ARACNE step enables *TGS-Lite+* to predict as many true positive edges as that of *TGS-Lite* while rejecting more number of false positive edges. That is why the former algorithm is able to retain the same recall as that of the latter algorithm while achieving a higher precision.

#### 4.11 Memory Usage: *TGS* vs *TGS-Lite*

This section presents the screenshots representing run-time memory usages of *TGS* and *TGS-Lite* for different values of the max fan-in ($M_f$) parameter.

```
top - 15:13:55 up 3 days, 2 min,  4 users,  load average: 1.37, 0.77, 1.01
Tasks:   1 total,   1 running,   0 sleeping,   0 stopped,   0 zombie
%Cpu(s):  6.7 us,  0.3 sy,  0.0 ni, 92.9 id,  0.0 wa,  0.0 hi,  0.1 si,  0.0 st
KiB Mem : 32933260 total, 24978164 free,  3309772 used,  4645324 buff/cache
KiB Swap: 35848188 total, 35848188 free,        0 used. 28883336 avail Mem

  PID USER      PR  NI    VIRT    RES    SHR S  %CPU %MEM     TIME+ COMMAND
30182 saptars+  20   0 1475992 1.222g  13720 R 100.0  3.9   1:35.46 R
```

(a) For $M_f = 20$, *TGS* uses 3.9% of memory (column '%MEM').

```
top - 15:32:27 up 3 days, 20 min,  4 users,  load average: 1.00, 1.22, 1.21
Tasks:   1 total,   1 running,   0 sleeping,   0 stopped,   0 zombie
%Cpu(s):  6.3 us,  0.5 sy,  0.0 ni, 93.2 id,  0.0 wa,  0.0 hi,  0.0 si,  0.0 st
KiB Mem : 32933260 total, 26037236 free,  2248104 used,  4647920 buff/cache
KiB Swap: 35848188 total, 35848188 free,        0 used. 29944228 avail Mem

  PID USER      PR  NI    VIRT    RES    SHR S  %CPU %MEM     TIME+ COMMAND
18605 saptars+  20   0  412452 217568  13752 R  99.7  0.7   1:36.57 R
```

(b) For $M_f = 20$, *TGS-Lite* uses 0.7% of memory (column '%MEM').

Figure 4.4: The Percentages of Memory Usage by *TGS* and *TGS-Lite* for $M_f = 20$. The dataset in use is Ds100n. The shown figures are screenshots of the 'top' command in the Ubuntu OS. These screenshots are taken when the Bene step begins which is around 1 minutes 30 seconds (column 'TIME+') after the execution starts.)

```
top - 15:21:56 up 3 days, 10 min,  4 users,  load average: 1.45, 1.08, 1.07
Tasks:   1 total,   1 running,   0 sleeping,   0 stopped,   0 zombie
%Cpu(s):  6.2 us,  0.3 sy,  0.0 ni, 93.4 id,  0.0 wa,  0.0 hi,  0.1 si,  0.0 st
KiB Mem : 32933260 total, 23794232 free,  4492420 used,  4646608 buff/cache
KiB Swap: 35848188 total, 35848188 free,        0 used. 27700192 avail Mem

  PID USER      PR  NI    VIRT    RES    SHR S  %CPU %MEM     TIME+ COMMAND
 7184 saptars+  20   0 2655760 2.347g  13564 R  99.7  7.5   1:37.89 R
```

(a) For $M_f = 21$, *TGS* uses 7.5% of memory (column '%MEM').

```
top - 15:36:27 up 3 days, 24 min,  4 users,  load average: 2.19, 1.63, 1.36
Tasks:   1 total,   1 running,   0 sleeping,   0 stopped,   0 zombie
%Cpu(s):  6.4 us,  0.4 sy,  0.0 ni, 93.0 id,  0.0 wa,  0.0 hi,  0.1 si,  0.0 st
KiB Mem : 32933260 total, 26037844 free,  2247216 used,  4648200 buff/cache
KiB Swap: 35848188 total, 35848188 free,        0 used. 29945272 avail Mem

  PID USER      PR  NI    VIRT    RES    SHR S  %CPU %MEM     TIME+ COMMAND
22829 saptars+  20   0  412452 217576  13760 R 100.0  0.7   1:42.09 R
```

(b) For $M_f = 21$, *TGS-Lite* uses 0.7% of memory (column '%MEM').

Figure 4.5: The Percentages of Memory Usage by *TGS* and *TGS-Lite* for $M_f = 21$. The dataset in use is Ds100n. The shown figures are screenshots of the 'top' command in the Ubuntu OS. These screenshots are taken when the Bene step begins which is around 1 minutes 30 seconds (column 'TIME+') after the execution starts.)

```
top - 15:24:22 up 3 days, 12 min,  4 users,  load average: 1.39, 1.12, 1.08
Tasks:   1 total,   1 running,   0 sleeping,   0 stopped,   0 zombie
%Cpu(s):  6.2 us,  0.3 sy,  0.0 ni, 93.5 id,  0.0 wa,  0.0 hi,  0.0 si,  0.0 st
KiB Mem : 32933260 total, 21318284 free,  6967460 used,  4647516 buff/cache
KiB Swap: 35848188 total, 35848188 free,        0 used. 25224576 avail Mem

  PID USER      PR  NI    VIRT    RES    SHR S  %CPU %MEM     TIME+ COMMAND
 9819 saptars+  20   0 5109452 4.687g  13452 R 100.0 14.9   1:37.52 R
```

(a) For $M_f = 22$, *TGS* uses 14.9% of memory (column '%MEM').

```
top - 15:39:13 up 3 days, 27 min,  4 users,  load average: 1.58, 1.77, 1.47
Tasks:   1 total,   1 running,   0 sleeping,   0 stopped,   0 zombie
%Cpu(s):  6.1 us,  0.3 sy,  0.0 ni, 93.5 id,  0.0 wa,  0.0 hi,  0.0 si,  0.0 st
KiB Mem : 32933260 total, 26030608 free,  2254108 used,  4648544 buff/cache
KiB Swap: 35848188 total, 35848188 free,        0 used. 29938308 avail Mem

  PID USER      PR  NI    VIRT    RES    SHR S  %CPU %MEM     TIME+ COMMAND
25885 saptars+  20   0  417248 222528  13820 R 100.0  0.7   1:37.29 R
```

(b) For $M_f = 22$, *TGS-Lite* uses 0.7% of memory (column '%MEM').

Figure 4.6: The Percentages of Memory Usage by *TGS* and *TGS-Lite* for $M_f = 22$. The dataset in use is Ds100n. The shown figures are screenshots of the 'top' command in the Ubuntu OS. These screenshots are taken when the Bene step begins which is around 1 minutes 30 seconds (column 'TIME+') after the execution starts.)

```
top - 15:27:33 up 3 days, 16 min,  4 users,  load average: 2.04, 1.57, 1.27
Tasks:   1 total,   1 running,   0 sleeping,   0 stopped,   0 zombie
%Cpu(s):  5.9 us,  0.9 sy,  0.0 ni, 93.2 id,  0.0 wa,  0.0 hi,  0.0 si,  0.0 st
KiB Mem : 32933260 total, 19499452 free,  8786200 used,  4647608 buff/cache
KiB Swap: 35848188 total, 35848188 free,        0 used. 23406024 avail Mem

  PID USER      PR  NI    VIRT    RES    SHR S  %CPU %MEM     TIME+ COMMAND
13276 saptars+  20   0 7137216 6.433g  13340 R 100.0 20.5   1:37.46 R
```

(a) For $M_f = 23$, *TGS* uses 20.5% of memory (column '%MEM').

```
top - 15:41:34 up 3 days, 30 min,  4 users,  load average: 1.05, 1.48, 1.40
Tasks:   1 total,   1 running,   0 sleeping,   0 stopped,   0 zombie
%Cpu(s):  6.3 us,  0.4 sy,  0.0 ni, 93.3 id,  0.0 wa,  0.0 hi,  0.0 si,  0.0 st
KiB Mem : 32933260 total, 26040940 free,  2243628 used,  4648692 buff/cache
KiB Swap: 35848188 total, 35848188 free,        0 used. 29948900 avail Mem

  PID USER      PR  NI    VIRT    RES    SHR S  %CPU %MEM     TIME+ COMMAND
28470 saptars+  20   0  412464 217492  13716 R 100.0  0.7   1:36.28 R
```

(b) For $M_f = 23$, *TGS-Lite* uses 0.7% of memory (column '%MEM').

Figure 4.7: The Percentages of Memory Usage by *TGS* and *TGS-Lite* for $M_f = 23$. The dataset in use is Ds100n. The shown figures are screenshots of the 'top' command in the Ubuntu OS. These screenshots are taken when the Bene step begins which is around 1 minutes 30 seconds (column 'TIME+') after the execution starts.)

```
top - 15:25:33 up 154 days, 22:34,  3 users,  load average: 2.08, 1.39, 1.37
Tasks:   2 total,   2 running,   0 sleeping,   0 stopped,   0 zombie
%Cpu(s):  9.2 us,  0.6 sy,  0.7 ni, 89.4 id,  0.0 wa,  0.0 hi,  0.1 si,  0.0 st
KiB Mem : 32933340 total,  2881680 free, 26610912 used,  3440748 buff/cache
KiB Swap: 35848188 total, 32574468 free,  3273720 used.  5457732 avail Mem

  PID USER      PR  NI    VIRT    RES    SHR S  %CPU %MEM     TIME+ COMMAND
27646 saptars+  20   0 17.994g 0.017t  13364 R 100.0 54.7   1:35.25 R
27694 saptars+  20   0  417432 222576  13688 R  99.7  0.7   1:27.44 R
```

Figure 4.8: The Percentages of Memory Usage (columns '%MEM') by *TGS* and *TGS-Lite* for $M_f = 24$. *TGS* uses 54.7% while *TGS-Lite* uses only 0.7%. The dataset in use is Ds100n. The shown figure is a screenshot of the 'top' command in the Ubuntu OS. This screenshot is taken when the Bene step begins which is around 1 minutes 30 seconds (column 'TIME+') after the executions start.)

#### 4.12 A Comparative Study with DmLc3 Sub-datasets

This section presents a comparative study between algorithms {*TGS-Lite.mf15, TGS-Lite+.mf15, ARTIVA, TVDBN-0, TVDBN-bino-hard, TVDBN-bino-soft*} with DmLc3 sub-datasets.

##### 4.12.1 Input and Output Files

The input and output files for {*TGS-Lite.mf15, TGS-Lite+.mf15*} are discussed in Chapter 3 . The input and output files for {*ARTIVA, TVDBN-0, TVDBN-bino-hard, TVDBN-bino-soft*} are given in Tables 4.2 , 4.3 , 4.4 , 4.5 , respectively.

Table 4.2: The Input JSON Files and Output Directories for the ARTIVA Algorithm. The input JSON files can be found at: `https://github.com/sap01/TGS-Lite-supplem/tree/master/input/ARTIVA` . The output directories can be found at: `https://github.com/sap01/TGS-Lite-supplem/tree/master/results/ARTIVA` . To run the ARTIVA code, please refer to Section 2.6 of the supplementary document of Pyne et al. [7].

| Sub-dataset | Input JSON File | Output Directory |
|---|---|---|
| DmLc3E | input.DmLc3E.json | ARTIVA-DmLc3E-output20190724180819 |
| DmLc3L | input.DmLc3L.json | ARTIVA-DmLc3L-output20190726145943 |
| DmLc3P | input.DmLc3P.json | ARTIVA-DmLc3P-output20190726155422 |
| DmLc3A | input.DmLc3A.json | ARTIVA-DmLc3A-output20190726153445 |

Table 4.3: The Input JSON Files and Output Directories for the TVDBN-0 Algorithm. The input JSON files can be found at: `https://github.com/sap01/TGS-Lite-supplem/tree/master/input/TVDBN-0` . The output directories can be found at: `https://github.com/sap01/TGS-Lite-supplem/tree/master/results/TVDBN-0` . To run the TVDBN-0 code, please refer to Section 2.7 of the supplementary document of Pyne et al. [7].

| Sub-dataset | Input JSON File | Output Directory |
|---|---|---|
| DmLc3E | input.DmLc3E.poisson.json | TVDBN-0-DmLc3E-output20190801141920 |
| DmLc3L | input.DmLc3L.poisson.json | TVDBN-0-DmLc3L-output20190801125954 |
| DmLc3P | input.DmLc3P.poisson.json | TVDBN-0-DmLc3P-output20190801131316 |
| DmLc3A | input.DmLc3A.poisson.json | TVDBN-0-DmLc3A-output20190801124701 |

Table 4.4: The Input JSON Files and Output Directories for the TVDBN-bino-hard Algorithm. The input JSON files can be found at: `https://github.com/sap01/TGS-Lite-supplem/tree/master/input/TVDBN-bino-hard` . The output directories can be found at: `https://github.com/sap01/TGS-Lite-supplem/tree/master/results/TVDBN-bino-hard` . To run the TVDBN-bino-hard code, please refer to Section 2.7 of the supplementary document of Pyne et al. [7].

| Sub-dataset | Input JSON File | Output Directory |
|---|---|---|
| DmLc3E | input.DmLc3E.bino_hard.json | TVDBN-bino-hard-DmLc3E-output20190731143633 |
| DmLc3L | input.DmLc3L.bino_hard.json | TVDBN-bino-hard-DmLc3L-output20190731141304 |
| DmLc3P | input.DmLc3P.bino_hard.json | TVDBN-bino-hard-DmLc3P-output20190731144806 |
| DmLc3A | input.DmLc3A.bino_hard.json | TVDBN-bino-hard-DmLc3A-output20190731140141 |

Table 4.5: The Input JSON Files and Output Directories for the TVDBN-bino-soft Algorithm. The input JSON files can be found at: `https://github.com/sap01/TGS-Lite-supplem/tree/master/input/TVDBN-bino-soft` . The output directories can be found at: `https://github.com/sap01/TGS-Lite-supplem/tree/master/results/TVDBN-bino-soft` . To run the TVDBN-bino-soft code, please refer to Section 2.7 of the supplementary document of Pyne et al. [7].

| **Sub-dataset** | **Input JSON File** | **Output Directory** |
|---|---|---|
| DmLc3E | input.DmLc3E.bino_soft.json | TVDBN-bino-soft-DmLc3E-output20190729170325 |
| DmLc3L | input.DmLc3L.bino_soft.json | TVDBN-bino-soft-DmLc3L-output20190731133429 |
| DmLc3P | input.DmLc3P.bino_soft.json | TVDBN-bino-soft-DmLc3P-output20190730153340 |
| DmLc3A | input.DmLc3A.bino_soft.json | TVDBN-bino-soft-DmLc3A-output20190729164910 |

***Note:*** The hardware configuration of the server used for running *TGS-Lite.mf15* and *TGS-Lite+.mf15* on DmLc3 sub-datasets is mentioned in Section 2.2 ('server 1'). Unfortunately, the server went out of service afterwards. Therefore, an equivalent server ('server 2') is deployed for running {*ARTIVA, TVDBN-0, TVDBN-bino-hard, TVDBN-bino-soft*} on DmLc3 sub-datasets. A parameter-wise comparison of these two servers are given in Table 4.6.

Table 4.6: A Comparison Between Two Computing Servers used for DmLc3 Sub-datasets

| **Parameter** | **Server 1** | **Server 2** |
|---|---|---|
| Architecture | x86 64 | x86 64 |
| CPU | Intel® Xeon® X5675 @ 3.07GHz | Intel® Xeon® W-2145 CPU @ 3.70GHz |
| Main Memory | 31 GB | 31 GB |
| Swap Space | 34 GB | 32 GB |
| Cache | {L1d cache: 32 KB, L1i cache: 32 KB, L2 cache: 256 KB, L3 cache: 12288 KB} | {L1d cache: 32 KB, L1i cache: 32 KB, L2 cache: 1 MB, L3 cache: 11264 KB} |
| Secondary Storage | 4.1 TB | 1 TB |
| Operating System | Ubuntu 12.04.5 LTS (Codename: Precise) | Ubuntu 16.04.6 LTS (Codename: Xenial) |

##### 4.12.2 Results

*TGS-Lite.mf15* and *TGS-Lite+.mf15* are able to process all sub-datasets. However, *ARTIVA* is only able to process the DmLc3E sub-dataset. For other sub-datasets, *ARTIVA* throws errors which can be found in the 'output.txt' files of the corresponding output directories. The authors are in the process of submitting a detailed bug report to the maintainer of the ARTIVA package.

*TVDBN-0*, *TVDBN-bino-hard* and *TVDBN-bino-soft* are unable to process any sub-dataset. They throw 'Cannot allocate memory' error as can be found out in the 'nohup...out' files inside the corresponding output directories. This is an interesting finding; the memory requirements of these algorithms are not discussed by Dondelinger et al. while proposing the algorithms [3]. Therefore, analysing their memory requirements presents a novel opportunity for future reviews.

In essence, only for the DmLc3E sub-dataset, the results are available for {*TGS-Lite.mf15*, *TGS-Lite+.mf15*, *ARTIVA*}. Hence, a comparative study is done between them following the evaluation strategy used in the main paper (Paragraph 'Evaluation Strategy', Section 5.2, the main paper). The complete set of findings can be found at:

https://github.com/sap01/TGS-Lite-supplem/blob/master/results/DmLc3.flybase.ids.redfly.tfs.xlsx. Some of these findings are discussed below.

***Gene 'Antp':*** Both *TGS-Lite.mf15* and *TGS-Lite+.mf15* predict 'opa' to be a regulatee of 'Antp' in DmLc3E. According to the existing biological knowledge, it is highly likely to be a true positive prediction (Paragraph 'Gene 'Antp'', Section 5.2, main paper). *ARTIVA* misses this edge. On the other hand, *TGS-Lite.mf15* and *TGS-Lite+.mf15* predict 17 and 9 other regulatees, respectively. There are hitherto no experimental evidences for these regulatees. *ARTIVA* rejects all of them which could be true negative predictions. Instead, *ARTIVA* predicts two regulatees of 'Antp' – 'hdc' and 'Sulf1'. For 'hdc', there is not sufficient information to validate its relationship with 'Antp'; please see: http://flybase.org/reports/FBgn0010113 . However, for 'Sulf1', there is sufficient information to argue that it is a false positive prediction. 'Sulf1' is found in the extracellular region whereas 'Antp' is found in nucleus and some other cellular components; please see Section 'GO Summary Ribbons' of 'Sulf1' ( http://flybase.org/reports/FBgn0040271 ) and 'Antp' ( http://flybase.org/reports/FBgn0260642 ). Therefore, they are less likely to have a regulatory relationship.

***Gene 'eve':*** *TGS-Lite.mf15* predicts three regulatees of 'eve' in DmLc3E – 'mmy', 'inv' and 'twi'. All three of them are potentially true positive predictions. The potential of 'mmy' is already discussed in Paragraph 'Gene 'eve'', Section 5.2 of the main paper. That of 'inv' can be argued based on the fact that both 'eve' and 'inv' are transcriptional repressors essential for segmentation in the embryo; please see Section 'Gene Snapshot' of 'eve' ( http://flybase.org/reports/FBgn0000606 ) and 'inv' ( http://flybase.org/reports/FBgn0001269 ). In case of 'twi', it is co-localised with 'eve' in nucleus; moreover, both of them take part in cell organisation/biogenesis; please see Section 'GO Summary Ribbons' of 'eve' ( http://flybase.org/reports/FBgn0000606 ) and 'twi' ( http://flybase.org/reports/FBgn0003900 ). Therefore, there is a high probability that they have a regulatory relationship. *TGS-Lite+.mf15* predicts 'twi' but misses 'inv' and 'mmy'. *ARTIVA*, on the other hand, misses all of them.

***Gene 'prd':*** *TGS-Lite.mf15* predicts 9 regulatees of 'prd' in DmLc3E. One of them is 'eve' which is a known regulatee. For other regulatees, hitherto there does not exist any experimental evidence. *TGS-Lite+.mf15* incorrectly rejects 'eve'. However, it also rejects 7 other regulatees, which are potentially true negative predictions. The only regulatee that *TGS-Lite+.mf15* predicts jointly with *TGS-Lite.mf15* is 'knrl'. It is also the only regulatee of 'prd' predicted by *TGS-Lite+.mf15*. 'Knrl' might be a true positive prediction given that it is co-localised with 'prd' in nucleus; please see Section 'GO Summary Ribbons' of 'prd' ( http://flybase.org/reports/FBgn0003145 ) and 'knrl' ( http://flybase.org/reports/FBgn0001323 ). On the other hand, *ARTIVA*'s predictions do not have any common regulatees with that of the aforementioned algorithms. Instead, *ARTIVA* predicts a single regulatee – 'LIMK1'. It is difficult to forecast the validity of this prediction given the current biological knowledge. In one hand, 'LIMK1' participates in reproduction whereas 'prd' has a known role in male fertility, which suggest that they might have a regulatory relationship; on the other hand, 'prd' is localised in nucleus but there is no evidence of 'LIMK1' being found in nucleus, which suggest that they are less likely to have a regulatory relationship; please see Sections 'Gene Snapshot' and 'GO Summary Ribbons' of 'prd' ( http://flybase.org/reports/FBgn0003145 ) and 'LIMK1' ( http://flybase.org/reports/FBgn0283712 ).

***Summary:*** The findings of this study can be summed up as follows.

- *Memory Management.* *TGS-Lite.mf15* and *TGS-Lite+.mf15* are able to process all DmLc3 sub-datasets with the given main memory (31 GB). *ARTIVA* is able to process the DmLc3E sub-dataset. It is not able to process other sub-datasets potentially due to some implementation issues, and not due to memory management issues. However, *TVDBN-0*, *TVDBN-bino-hard* and *TVDBN-bino-soft* fail to process all sub-datasets due to memory insufficiency. As a result, the comparative study is performed between the results of *TGS-Lite.mf15*, *TGS-Lite+.mf15* and *ARTIVA* with the DmLc3E sub-dataset.

- *Learning Power.* *ARTIVA* makes less number of potentially false positive predictions than that of *TGS-Lite.mf15*; however, the latter algorithm makes more number of potentially true positive predictions. *TGS-Lite+.mf15*, provides a balance between the aforementioned algorithms. It makes less number of potentially false positive predictions than that of *TGS-Lite.mf15*; at the same time, it produces more number of potentially true positive predictions than that of *ARTIVA*. Therefore, the comparative reconstruction powers of these algorithms on the real dataset DmLc3E is consistent with that on the in-silico datasets (Section 5.1 of the main paper).

- *Learning Speed.* The same consistency is preserved with their comparative learning speeds. *ARTIVA* remains the slowest, taking around 7 hours for DmLc3E. *TGS-Lite.mf15* performs relatively faster, clocking around 4 hours. *TGS-Lite+.mf15* again achieves the fastest performance, taking only about 22 minutes.

##### 4.12.3 Shortcut for Verifying Results

For readers' convenience, a shortcut method is designed for verifying the results of the comparative study. For example, if the reader wishes to verify the results presented in Paragraph 'Gene 'Antp'' in Section 4.12.2 , then he/she needs to know the names of the regulatees of 'Antp' predicted by *TGS-Lite.mf15 / TGS-Lite+.mf15 / ARTIVA* for the DmLc3E sub-dataset. The user can find out the names using the following procedure:

```
%% Download the R session file:
%% https://github.com/sap01/TGS-Lite-supplem/blob/master/results/sess.eval.mf15.RData

$$ Then load it into an R session.
> load('sess.eval.mf15.RData')

%% Print the names of the regulatees of 'Antp' predicted by TGS-Lite.mf15 from DmLc3E
> names(which(TGS.Lite.mf15.di.net.adj.matrix.E["Antp", ] == 1))
 [1] "btl"          "siz"          "cp309"        "Atg6"         "FKBP59"
 [6] "G.salpha60A" "rib"          "run"          "pnt"          "exu"
 [11] "odd"          "bl"           "alpha.Spec"  "disco"        "opa"
 [16] "stil"         "aft"          "CG12896"

%% Print the names of the regulatees of 'Antp' predicted by TGS-Lite+.mf15 from DmLc3E
> names(which(TGS.Lite.plus.mf15.di.net.adj.matrix.E["Antp", ] == 1))
 [1] "pros"        "btl"         "siz"         "rib"         "run"
 [6] "pnt"         "exu"         "alpha.Spec" "opa"         "CG12896"

%% Print the names of the regulatees of 'Antp' predicted by ARTIVA from DmLc3E
> names(which(ARTIVA.di.net.adj.matrix.E["Antp", ] == 1))
[1] "hdc"   "Sulf1"

%% Similarly, to print the names of the regulatees of some other gene,
%% replace 'Antp' with the desired gene's name.
```
